# Cancer-associated fibroblasts serve as decoys to suppress NK cell anti-cancer cytotoxicity

**DOI:** 10.1101/2023.11.23.568355

**Authors:** Aviad Ben-Shmuel, Yael Gruper, Oshrat Levi-Galibov, Hallel Rosenberg-Fogler, Giulia Carradori, Yaniv Stein, Maya Dadiani, Mariia Naumova, Reinat Nevo, Dana Morzaev-Sulzbach, Gal Yagel, Shimrit Mayer, Einav Nili Gal-Yam, Ruth Scherz-Shouval

**Affiliations:** Department of Biomolecular Sciences, The Weizmann Institute of Science, Rehovot; Chaim Sheba Medical Center, Cancer Research Center, Tel-Hashomer, Israel; Chaim Sheba Medical Center, Institute of Oncology, Tel-Hashomer, Israel

## Abstract

Cancer associated fibroblasts (CAFs) are among the most abundant components of the breast tumor microenvironment (TME) and major contributors to immune modulation. CAFs are well-known to regulate the activity of diverse types of immune cells including T cells, macrophages and dendritic cells, however little is known about their interaction with Natural killer (NK) cells, which constitute an important arm of anti-tumor immunity. Here we find, using mouse models of cancer and ex-vivo co-cultures, that CAFs inhibit NK cell cytotoxicity towards cancer cells. We unravel the mechanism by which this suppression occurs, through ligand-receptor engagement between NK cells and CAFs leading to CAF cytolysis, which in turn diminishes the expression of activating receptors on NK cells, promoting cancer escape from NK cell surveillance. Analysis of breast cancer patient samples reveals enrichment of NK cells in CAF-rich regions, and upregulation of NK binding ligands on CAFs which is correlated with poor disease outcome. These results reveal a CAF-mediated immunosuppressive decoy mechanism with implications for treatment of solid tumors.

## Introduction

Cancer associated fibroblasts (CAFs) are among the most abundant components of the tumor microenvironment (TME) in carcinomas and major protumorigenic players ^1^. CAFs promote cancer through secretion of extracellular matrix components, growth factors and cytokines that reshape the composition of the TME ^1–3^. In recent years it has become evident that CAFs play a central role in regulating immune responses in the TME^4^. However most studies focused on their ability to modulate T-cells and macrophages, and much less is known regarding their cross-talk with other components of the immune system.

CAFs are composed of heterogeneous subpopulations proposed to originate from different cell types including tissue resident fibroblasts, mesenchymal bone marrow cells, adipocyte-derived precursor cells, endothelial cells, mesothelial cells, and pericytes^5–11^. CAF subpopulations are also heterogeneous and plastic in their phenotypes and tasks, and can adopt contextual functions, depending on the cancer subtype, mutational status of the cancer cells, and specific organ in which the cancer develops^5,12–18^.

While traditionally viewed as wound healing matrix remodeling cells, we know today that CAFs are major modulators of the immune TME^4,19^ and hamper the efficacy of immunotherapy^20–23^. Distinct mechanisms that induce immune dysfunction have been attributed to CAFs, most of them affecting T cells and macrophages. In the context of T-cell immunity, CAFs mediate suppression through 3 modalities^4^: CD8^+^ T-cell exclusion^24^, impairment of CD4^+^/CD8^+^ activation and persistence^25,26^, and recruitment and expansion of suppressive T-regulatory (Treg) cells^27,28^. CAF-mediated regulation of macrophages occurs mainly through recruitment of monocytes and induction of the pro-tumorigenic tumor associated macrophage (TAM) phenotype^4^ fueled by reciprocal paracrine interactions^29–33^. Additional work has been conducted on CAF regulation of other immune cell populations including neutrophils^34,35^ and dendritic cells (DCs)^28,36^.

One of the understudied immune populations potentially affected by CAFs is natural killer (NK) cells that play an important role in cancer mitigation. NK cells are a principal component of the innate immune system. They directly target virally infected cells and cancer cells through secretion of cytolytic granules^37^. In addition, they orchestrate adaptive immunity through secretion of cytokines such as interferon gamma (IFNγ), and stimulation of DC recruitment and activation through secretion of XCL1/2 and FLT3LG^38^. NK cell recognition and elimination of target cancer cells is mediated via the combined activation and inhibition of a variety of cell surface receptors. Activating receptors act via recognition of ligands that are upregulated on transformed and virally infected cells. Principal inhibitory receptors prevent NK cells from damaging or killing healthy ‘self’ cells through recognition of major histocompatibility class I (MHC-I), which is often downregulated on cancer cells as an immune escape mechanism^39^. Additional inhibitory checkpoint receptors are expressed on the surface of NK cells, and the delicate balance of signaling inputs from stimulatory and inhibitory receptors ultimately dictates the formation of an activating NK immune synapse and cytotoxicity against the target cell^40–42^.

The ability of NK cells to identify target cells without the requirement of antigen specificity, and their relatively low toxicity profile in clinical trials, makes them attractive candidates for immunotherapy^43^. Nevertheless, their capacity to persist and invoke anti-tumor immunity is limited in advanced solid tumors, due to different restraining responses^44^. Despite the high abundance of CAFs in solid tumors, little is known regarding their possible effects on NK cell tumor surveillance, and mechanistic insights are lacking. Recent evidence in pancreatic ductal adenocarcinoma (PDAC) points to a Nertin G1 expressing CAF subset that inhibits NK cell cytotoxicity and cytokine production, partially through secretion of inhibitory cytokines and partially through downregulation of the pleiotropic Il-15 cytokine^45^. In melanoma, co-culture of allogeneic NK donor cells with established melanoma cell lines reduced NK cell cytotoxicity, primarily through a Prostaglandin E2 (PGE2) mediated mechanism facilitated through CAFs^46^. CAFs were also shown to indirectly influence NK cell function through recruitment and polarization of tumor associated macrophages^47^. While these studies point to potential regulation of NK cell activity by CAFs, we lack mechanistic studies of NK-CAF interactions, in particular studies performed under matched, autologous MHC-I systems. Furthermore, though CAFs heavily infiltrate breast tumors and contribute immensely to breast cancer pathology^2, 6,7, 14, 18,48^, no studies have deeply investigated the potential regulation of NK cells by CAFs in this disease. In breast cancer, multiple lines of evidence point to better outcomes for patients with tumors infiltrated by functional NK cells, and to an association between decreased NK cell function and breast cancer progression in patients and mouse models^49–53^. Therefore, understanding if and how NK cells are regulated by CAFs is pertinent to efforts to advance treatment strategies for breast cancer.

Here we address this question by examining NK-CAF-cancer interactions in two mouse models of triple negative breast cancer (TNBC). We employ these models to isolate CAFs from tumors and assess their ability to regulate matched NK cell activity in an *ex-vivo* co-culture setting, and find that NK cell cytotoxicity against cancer cells is reduced following physical interactions with CAFs. Mechanistic analysis revealed potent downregulation of the activating receptors DNAX Accessory Molecule-1 (DNAM-1) and Natural Killer Group 2D (NKG2D). Antibody blockade of NKG2D and DNAM-1 showed their necessity for NK cell elimination of TNBC cells in both models. Intriguingly, we found that CAFs isolated from tumors upregulate ligands for DNAM-1 and NKG2D, which are usually overexpressed on cancer cells and largely absent on healthy tissue, sensitizing CAFs to NK cell-mediated cytolysis. Indeed, CAFs were more sensitive to NK-mediated cytotoxicity than cancer cells in tri-culture experiments. Decoupling NK-CAF interactions with antibody blockade of DNAM-1 and NKG2D ligands resulted in partial rescue of DNAM-1 and NKG2D expression on NK cells and higher cytotoxicity against cancer cells. Analysis of human TNBC samples revealed enrichment of NK cells in stroma-rich vs. cancer-rich areas, and an association between high stromal expression of the DNAM-1 ligand, NECTIN2, and poor disease outcome. Together, these findings suggest a ‘decoy’ mechanism initiated in the tumor stroma that may have implications for immune-based therapies.

## Results

### CAFs impair NK cell cytotoxicity against cancer cells

To begin to investigate the potential regulation of NK cells by CAFs, we first asked whether CAFs affect the ability of NK cells to kill cancer cells. To that end, we designed a two-step *ex-vivo* co-culture assay of syngeneic primary NK cells, CAFs and cancer cells, using the 4T1 TNBC model. We previously mapped the single-cell CAF landscape of TNBC, revealing two major subpopulations of CAFs present in tumors which we termed sCAFs and pCAFs, based on hallmark expression of the genes *S100a4* and *Pdpn*, respectively^14^. To test the effect of breast CAFs on NK cell cytotoxicity, we isolated pCAFs and sCAFs from 4T1-GFP tumor-bearing mice^14^, and primary NK cells (pNKs) and normal mammary fibroblasts (NMFs) from syngeneic naive BALB/c mice (Figure 1A and Supplementary Figures 1A-B). We then performed a two-step co-culture assay: in the first step we activated the isolated pNKs with IL-2 and IL-15 in the presence or absence of the three fibroblast subsets for 24 hours. In the second step, we co-cultured the preconditioned NK cells with 4T1 cancer cells and assessed their cytotoxic activity. Co-culture with NMFs or sCAFs had no effect on pNK cell activity (Figure 1B). In stark contrast, co-culture of pNK cells with pCAFs significantly inhibited their cytotoxic capacity against 4T1 cells (Figure 1B), strengthening our previous observation of pCAF suppression of CD8^+^ T-cells in the 4T1 model^14^. The inhibitory activity of pCAFs was lost when conditioned medium (CM) from pCAFs was used instead of direct co-culture, suggesting that CAF-mediated inhibition of NK cytotoxicity against cancer cells requires direct interactions between CAFs and NK cells (Figure 1C).

**Figure 1.**
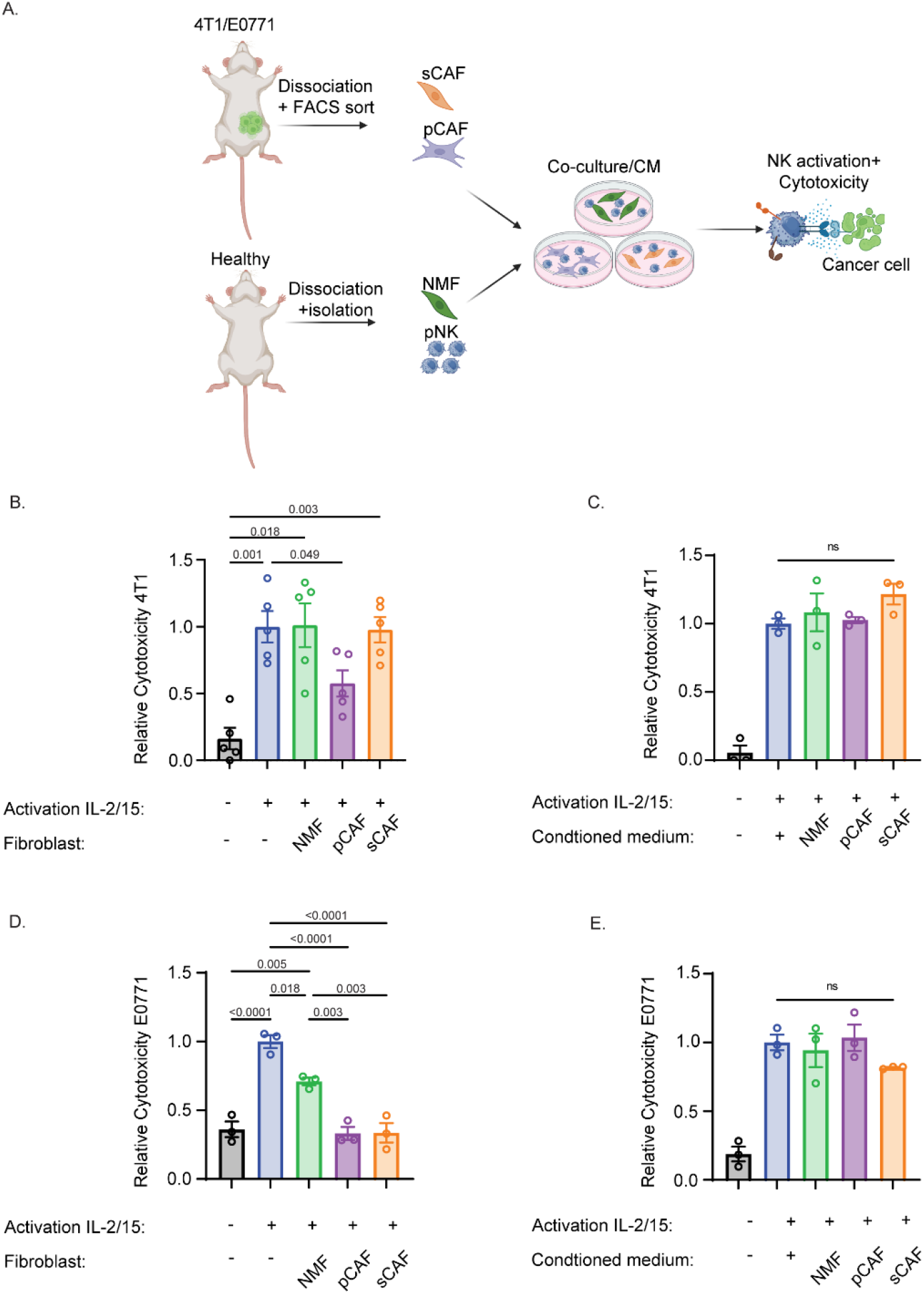
CAFs impair the cytotoxic capacity of NK cells towards cancer cells. (A) Schematic representation of the workflow. CAFs isolated from 4T1/E0771 tumors or NMFs isolated from healthy syngeneically matched mice were co-cultured with NK cells in the presence of the stimulating cytokines IL-2 and IL-15 for 24 hours. Subsequently, NK cells were removed and their cytotoxicity against matched 4T1/E0771-Luc cells was assessed by measurement of the luminescence of cancer cells. The image was generated using www.Biorender.com (**B, D**) NK cells from BALB/C (B) or C57BL/6 (D) mice were activated in the presence of each fibroblast subset for 24 hours, and then incubated with 4T1 (B) or E0771 (D) cells at a 5:1 NK:cancer ratio to determine cytotoxicity. The relative cytotoxicity (normalized to cytotoxicity of NK cells activated alone) is presented. N=6 biologically independent experiments for B, N=3 biologically independent experiments for D; data is presented as mean ± SEM. (**C, E**) NK cells from BALB/C (C) or C57BL/6 (E) mice were activated in RPMI or conditioned media from each fibroblast subset for 24 hours, and then incubated with 4T1 (B) or E0771 (D) cancer cells at a 5:1 NK:cancer ratio to determine cytotoxicity. The relative cytotoxicity (normalized to NK cells activated in RPMI) is presented. N=3 biologically independent experiments, data is presented as mean ± SEM.

These findings are not limited to the 4T1 model, as we observed a similar inhibitory effect of CAFs isolated from E0771 TNBC tumors in C57BL/6 mice (Figure 1D and Supplementary Figure 2A). As in the 4T1 model, activation of pNKs in the presence of CM from all fibroblast subpopulations did not alter their cytotoxic capacity against syngeneic E0771 cancer cells (Figure 1E). Co-culture with NMFs resulted in a partial reduction in NK cell cytotoxicity, while in this model, preincubation with either pCAFs or sCAFs resulted in a significantly greater reduction in NK cell cytotoxicity against E0771 cells (Figure 1D). Collectively these results suggest that direct interactions between CAFs and NK cells abrogate NK cell cytotoxic activity towards cancer cells.

### Surface expression of NKG2D and DNAM-1 on NK cells is impaired following interactions with CAFs

NK cells express a complex array of activating and inhibitory receptors, whose relative balance dictates cytolytic activity^57^. We hypothesized that CAFs may exert their inhibitory effect on NK cells by affecting the expression of these receptors. To test this, we monitored the cell-surface expression of a panel of activating and inhibitory receptors on pNK cells following *in-vitro* activation alone or in the presence of matching CAFs. Multiplexed flow cytometry analysis showed no significant changes in the expression of inhibitory/checkpoint receptors (LY49a, TIGIT, PD-1, KLRG1, LAG3, TIM-3) between NK cells activated alone or in the presence of fibroblasts (Supplementary Figure 2B and Supplementary Figure 3A-F). The expression of key activating receptors, however, was significantly affected by co-culture with fibroblasts: NKp46 and 2B4 were partially decreased, and the surface expression of NKG2D and DNAM-1 was almost completely abolished, diminishing to levels on-par or lower than unstimulated quiescent NK cells (Figure 2A-B and Supplementary Figure 3G-J).

**Figure 2.**
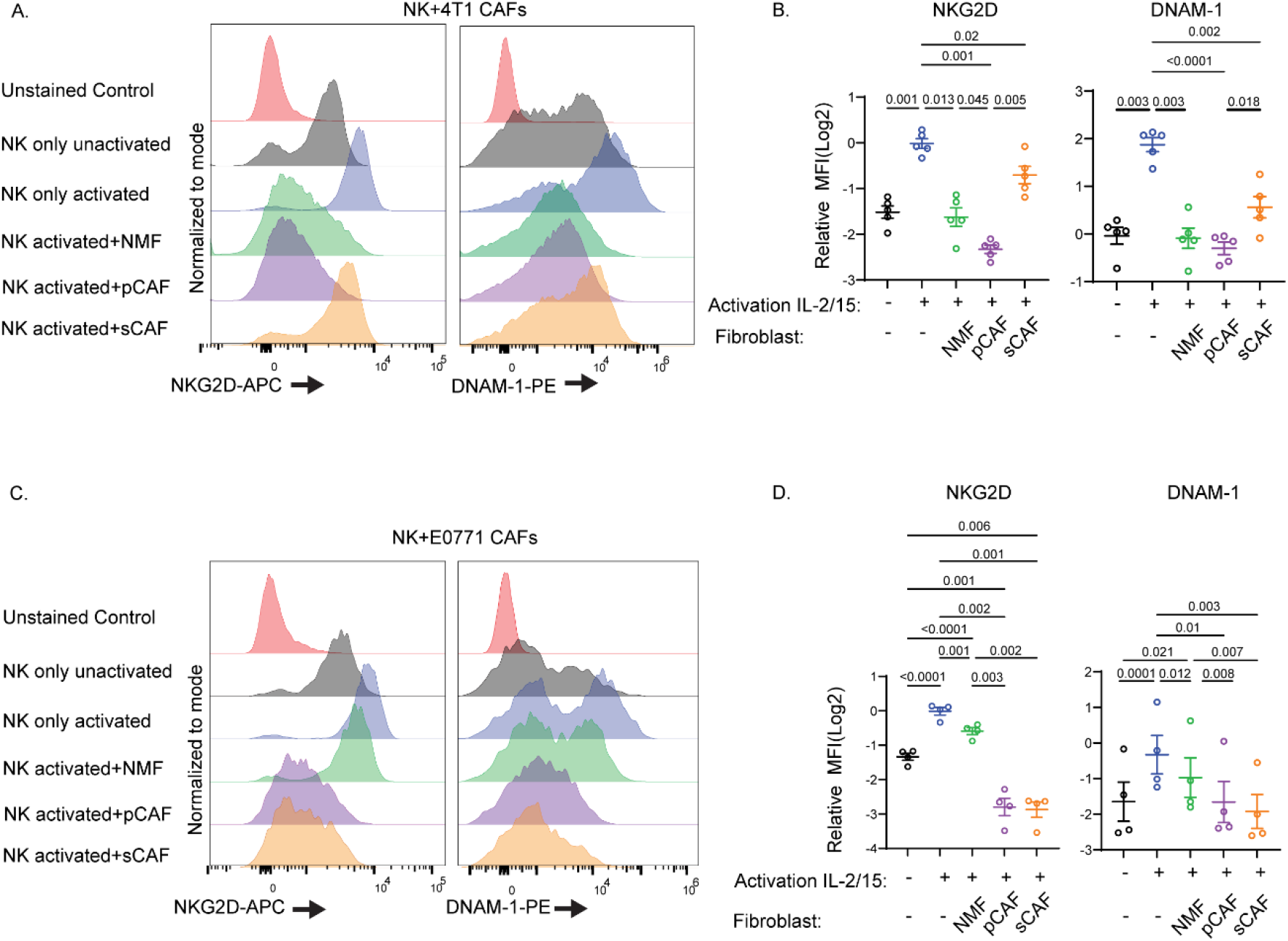
Co-culture with CAFs leads to inhibition of NKG2D and DNAM-1 expression on NK cells. NK cells from BALB/C **(A-B)** or C57Bl/6 mice **(C-D)** were activated with IL-2/15 alone or in the presence of different fibroblast subsets (CAFs isolated from 4T1 (A-B) or E0771 (C-D) tumors and NMFs isolated from healthy matching controls) for 24 hours. (**A, C**) Representative FACS plots of NKG2D and DNAM-1 expression on NK cells. The histograms presented are normalized to the modal value. (**B, D**) Quantification of the FACS experiments. N=5 biologically independent experiments for (B) and N=4 biologically independent experiments for (D), data is presented as mean ± SEM relative to NK cells activated alone.

Similar to the cytotoxicity experiments, fibroblast CM presented a marginal reduction in NKG2D and DNAM-1 expression compared to NK cells activated with control medium (Supplementary Figure 4A), further demonstrating the necessity of direct interactions for CAFs to mediate NK cell dysfunction.

To validate that the CAF-induced modulation of NKG2D and DNAM-1 expression on NK cells also occurs in other breast cancer models, we conducted orthogonal staining experiments using the E0771 tumor model. Similar to the 4T1 model, pNK cells cultured with CAFs derived from E0771 tumors displayed near complete abolishment of NKG2D and DNAM-1 receptor expression. Unlike the 4T1 model, co-culture of pNK cells with NMFs in this model modestly affected receptor expression (Figure 2C-D). In addition, unlike the 4T1 model, CM from fibroblasts in the E0771 model did result in reduction of both NKG2D and DNAM-1 expression on NK cells, albeit to a lesser extent, suggesting that in this model there may be additional secreted factors regulating surface expression of these receptors (Supplementary Figure 4B).

Together these data indicate that direct interactions with CAFs promote potent reduction of activating receptor expression on NK cells, potentially driving the observed inhibition of cytotoxicity.

### The cytotoxic activity of NK cells towards cancer cells is mediated through NKG2D and DNAM-1

Ligands for NKG2D and DNAM-1 are often upregulated during inflammation, stress, and cancer^54,55^ and could potentially mediate the killing by NK cells in breast cancer. Indeed, FACS analysis showed that both 4T1 and E0771 cells express the DNAM-1 ligands PVR/CD155 and Nectin2 and the NKG2D ligand RAE1 (4T1 Figure 3A-C, E0771 Figure 3F-H, and Supplementary Figure 5A-B). 4T1 cells (but not E0771 cells) also express the NKG2D ligands H60a^56^ and MULT1 (4T1 Figure 3D-E, E0771 Figure 3I). The finding that the two cancer cell lines express ligands for NKG2D and DNAM-1 suggests that these receptors could be mediating the killing of the cancer cells, and that their downregulation may play critical roles in TNBC surveillance by NK cells.

**Figure 3.**
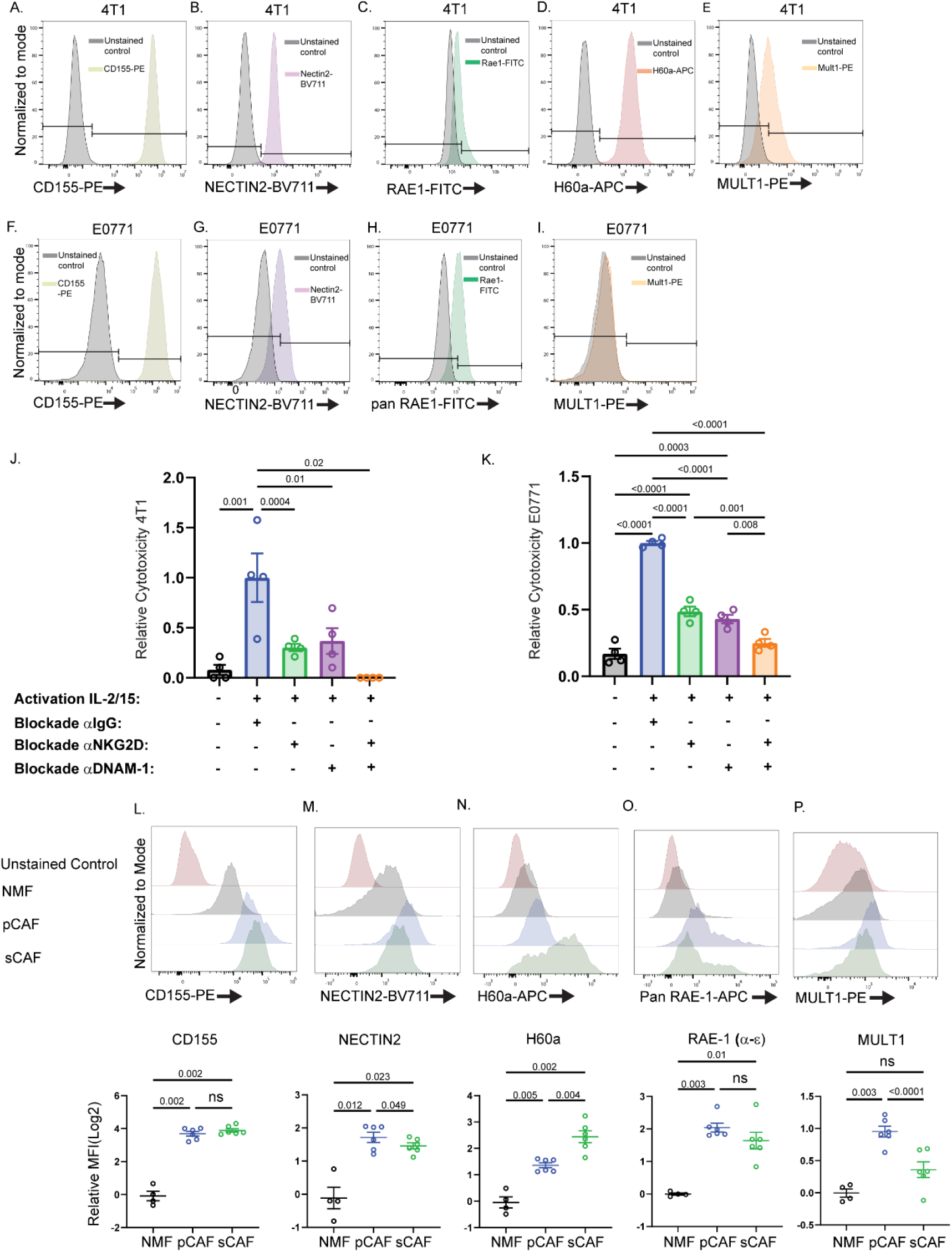
NKG2D and DNAM-1 regulate the recognition and cytotoxicity of TNBC cells. (A-I) FACS staining of 4T1 and E0771 TNBC cells for ligands of the NKG2D and DNAM-1 NK cell receptors (H60a is not stained for in E0771 cells as it is not expressed in the C57BL/6 background). Histograms presented are normalized to the modal value. (J-K) NK cells from BALB/c or C57BL/6 mice were activated for 24 hours, and then incubated with 4T1 or E0771 cells at a 5:1 NK:cancer ratio to determine cytotoxicity. NK cells were treated with either 50μg/ml IgG control or combinations of 50μg/ml α-NKG2D/DNAM-1 antibodies, as indicated. The relative cytotoxicity (normalized to cytotoxicity of NK cells activated with IgG control) is presented. N=4 biologically independent experiments for A and B, data is presented as mean ± SEM. (L-P) NMFs, pCAFs, and sCAFs freshly isolated from normal mammary fat pads of naive BALB/c mice or 4T1 tumors were subjected to FACS staining for the indicated NKG2D and DNAM-1 ligands. Quantification of the FACS experiments is shown in the bottom panels. N=4 mice for NMFs and N=6 mice for CAFs from 4T1 tumors. Data is normalized to stained NMFs samples, log2 transformed, and presented as mean ± SEM. The representative histograms (bottom) were normalized to the modal value.

To directly test this, we blocked each of the two receptors - NKG2D and DNAM-1 - separately or in combination, using blocking antibodies, and monitored the effect of this blockade on NK cytotoxicity against 4T1 or E0771 cells. Blockade of either NKG2D or DNAM-1 severely impaired NK cell cytotoxicity against both cancer cell lines, and dual blockade led to a synergistic and enhanced inhibitory effect on cytotoxicity (Figure 3J-K). Taken together these data demonstrate that TNBC cells express ligands for the NKG2D and DNAM-1 receptors present on activated NK cells, which mediate their effector activity. Downregulation of these receptors on NK cells by CAFs may therefore serve as a mechanism mediating TNBC escape from immune surveillance.

### CAFs upregulate surface expression of ligands for DNAM-1 and NKG2D

Surface expression of NKG2D and DNAM-1 could be regulated at the transcriptional level or post transcriptionally, whereby receptors are internalized and degraded following ligand binding^57–63^. To test whether ligands for NKG2D and DNAM-1 are also expressed on CAFs, we first leveraged bulk RNA-seq data that we previously acquired of NMFs and CAFs from 4T1 tumors^2,30^. We found that *Pvr* and *Nectin2*, encoding the ligands for DNAM-1, were upregulated in pCAFs and sCAFs compared to NMFs (Supplementary Figure 6A and Supplementary Table 1). In fact, *Nectin2* was among the most highly differentially expressed genes between pCAFs and NMFs, and was also significantly upregulated in pCAFs vs. sCAFs. The ligands for NKG2D were scarcely detected in all groups and their transcriptional regulation could therefore not be assessed.

To directly monitor the cell surface protein expression of NKG2D and DNAM-1 ligands on NMFs and CAFs, we isolated cells from normal mammary fat pads or 4T1 and E0771 tumors and stained for the ligands’ protein expression (Figure 3L-P and Supplementary Figure 6B-C). In both models the cell-surface expression of CD155 and Nectin2 was higher on both CAF subsets compared to NMFs (Figure 3L-M and Supplementary Figure 6C). In addition, higher expression of most NKG2D ligands was observed on CAFs compared to NMFs (Figure 3N-P and Supplementary Figure 6C).

Fibroblasts are characterized by high plasticity, and are sensitive to tuning by properties of the environment. Culturing on stiff 2D substrates was shown to induce activation in quiescent fibroblasts, reminiscent of a CAF-like state with high expression of fibroblast activation protein (FAP) and alpha-smooth muscle actin (αSMA)^64–66^. Therefore, we also assessed ligand expression on NMFs and CAFs following 4 days of 2D *in-vitro* culture (Supplementary Figure 7A-B). We found that the expression of CD155, NECTIN2, RAE1, and MULT1 on NMFs was highly induced following 4 days of culture. These results are concordant with the observation that cultured NMFs can acquire “CAF like” phenotypes and induce partial reduction in expression of NK cytotoxicity and NKG2D and DNAM-1 expression (Figures 1 and 2). They are also consistent with our prior observations that NMFs are the precursors of pCAFs in TNBC, were their rewiring and activation promote tumor growth and immune supression^14^. Collectively these data indicate that in the context of cancer development and progression, CAFs upregulate surface expression of ligands for DNAM-1 and NKG2D, possibly affecting NK cell receptor expression and activity.

### NK cells can kill CAFs via NKG2D and DNAM-1 engagement

NK cells were shown to interact with stromal cells and exert cytotoxic activity towards them under different pathological conditions, including hepatic fibrosis and rheumatoid arthritis. In both conditions, cell killing is mediated through NKG2D^67^ and DNAM-1^68^ ligand-receptor engagement. Furthermore, a previous study in melanoma demonstrated that downregulation of DNAM-1 required direct NK:CAF cell-cell contact, yet the precise mechanism was not elucidated^46^. We thus hypothesized that activation of quiescent fibroblasts and transition towards a CAF identity may induce NK cell cytotoxicity towards CAFs, leading to downregulation of NKG2D and DNAM-1 on NK-cells.

To test whether NK cells can kill fibroblasts, we measured the direct cytotoxicity of NK cells against NMFs, pCAFs, and sCAFs following 4 days of recovery after isolation. All fibroblast subsets were highly sensitive to NK cell-mediated cytotoxicity (Figure 4A-B), that was dramatically reduced following blockade with anti-NKG2D and anti-DNAM1 antibodies, suggesting that, similar to cancer cells, the killing of fibroblasts by NK cells is mediated through NKG2D and DNAM-1. These findings further suggest that CAFs may serve as “decoys” that engage NK cells and induce the subsequent downregulation of their activating receptors, thus preventing the killing of cancer cells.

**Figure 4.**
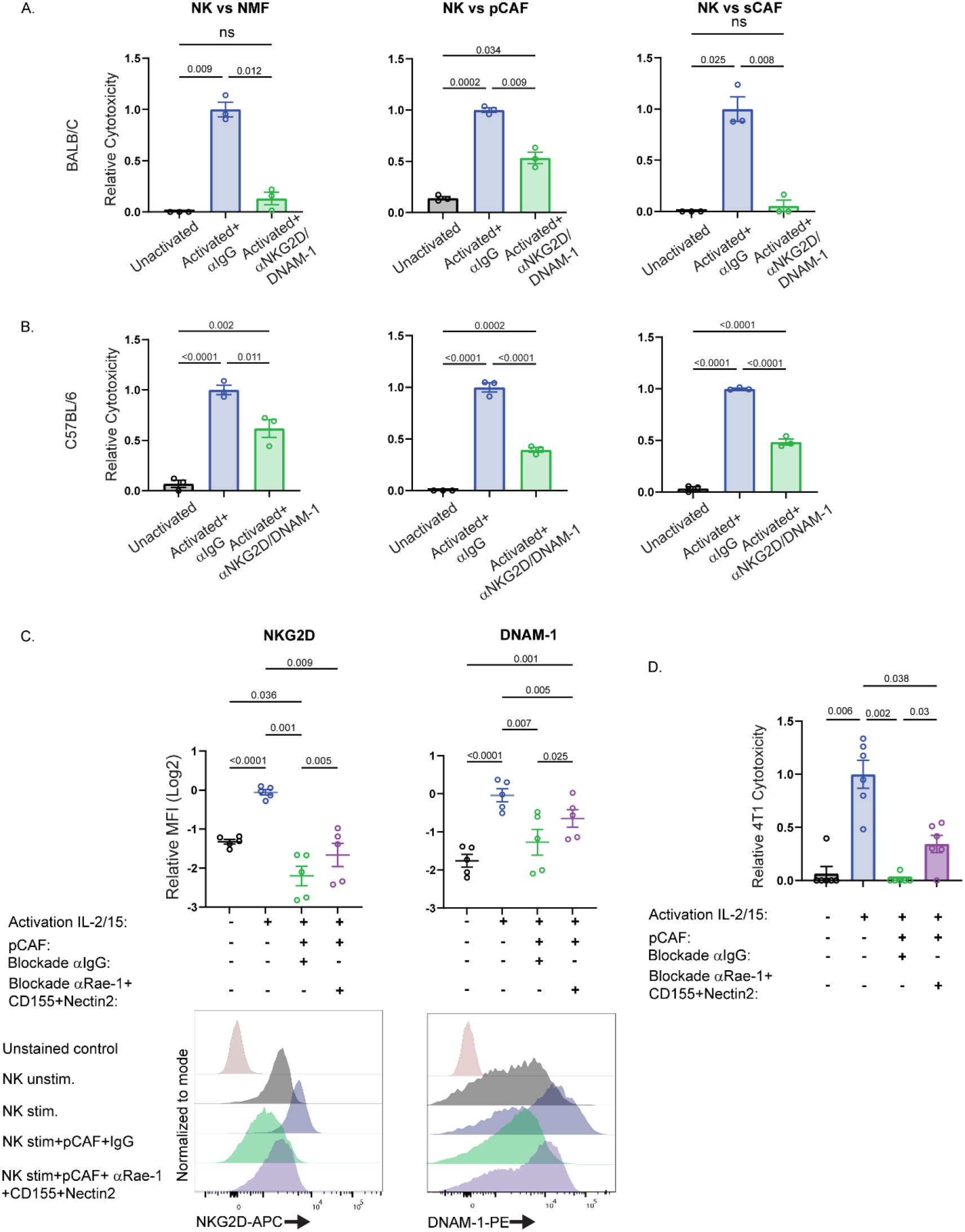
Decoupling of NKG2D/DNAM-1 interactions between CAFs and NK cells partially restores receptor expression and NK cell cytotoxicity. (A-B) NK cells from BALB/c or C57BL/6 mice were activated for 24 hours, and then incubated with NMFs, pCAFs, or sCAFs at a 5:1 NK:fibroblast ratio to determine cytotoxicity. NK cells were treated with either 50μg/ml IgG control or combinations of 50μg/ml α-NKG2D/DNAM1 antibodies to determine dependency on either receptor for cytotoxicity of fibroblast targets. The relative cytotoxicity (normalized to cytotoxicity of NK cells activated with IgG control) is presented (N=3 biologically independent experiments, data is presented as mean ± SEM). (C) NK cells from BALB/c mice were activated for 24 hours alone or in the presence of pCAFs that were pre-incubated with either 50μg/ml IgG control or combinations of 50μg/ml α-CD155, Nectin2, and Rae-1 antibodies. Subsequently, NK cells were subjected to FACS staining for NKG2D and DNAM-1. Quantification of the FACS experiments is shown in the top panel. N=5 biologically independent experiments, data is presented as mean ± SEM. The representative histograms (bottom) were normalized to the modal value. (D) NK cells from BALB/c mice were activated for 24 hours alone or as in (C), and then incubated with 4T1 cancer cells at a 5:1 NK:cancer ratio to determine cytotoxicity. Cytotoxicity (normalized to cytotoxicity of NK cells activated alone) is presented. N=6 biologically independent experiments, data is presented as mean ± SEM.

To test this hypothesis, we pretreated CAFs with blocking antibodies against pan-RAE-1, NECTIN-2, and CD155 (based on the highest expression of these markers on pCAFs relative to NMFs) or control IgG, followed by incubation with NK cells. We then assessed NK cell surface expression of NKG2D and DNAM-1, and cancer killing ability (Figure 4C-D). Activation of NK cells in the presence of blocked pCAFs resulted in partial rescue of NKG2D and DNAM-1 expression on NK cells and, importantly, partially rescued NK cell cytotoxicity against 4T1 cells (Figure 4D). These results indicate that NK cell-specific cytotoxicity against CAFs may lead to decreased expression of NKG2D and DNAM-1 receptors, which in turn impairs their ability to kill cancer cells in their surroundings.

### CAFs can compete with cancer cells for NK cell cytotoxicity and display greater sensitivity to NK-mediated lysis

Our finding that NK cells can kill CAFs suggests that CAFs compete with cancer cells as targets for NK cell killing. But are they equally sensitive to NK cell killing? To test this, we performed live cell imaging and quantitative analysis of NK cell killing in tri-cultures with CAFs and 4T1 cancer cells. We used the retraction of cell area^69^ as a proxy to evaluate cell death^70^, and cytopainter far-red and GFP to mark CAFs and 4T1 cells, respectively (Figure 5A-C, Supplementary Figure 8A-C and Supplementary Videos 1-5). Both CAFs and 4T1 cells were susceptible to NK-mediated cytotoxicity, supporting our previous cytotoxicity and receptor blockade assays (Figure 3J-K and Figure 4A-B). Intriguingly however, CAFs appeared to be much more sensitive to NK-mediated lysis compared to 4T1 cancer cells, dying at faster kinetics (Figure 5A-B and Supplementary Videos 1-5). Indeed, quantification of NK cell-mediated cytotoxicity demonstrated that CAFs were significantly more sensitive to NK cell cytolysis compared to 4T1 cells (Figure 5C). Moreover, at timepoints where all CAFs in a given field were eliminated, residual 4T1 cells still remained. Superior NK cytotoxic kinetics of CAFs were evident both in tri-culture conditions (CAFs with 4T1 cells and NK cells) and co-culture conditions (CAFs with NK cells only) (Supplementary Figure 8B-C). Finally, cytotoxicity experiments of activated NK cells seeded onto 4T1-luc cells alone (co-culture) or onto 4T1-luc cells and pCAFs (tri-culture) revealed significantly lowered cytotoxicity against cancer cells when CAFs were present in culture (Figure 5D).

**Figure 5.**
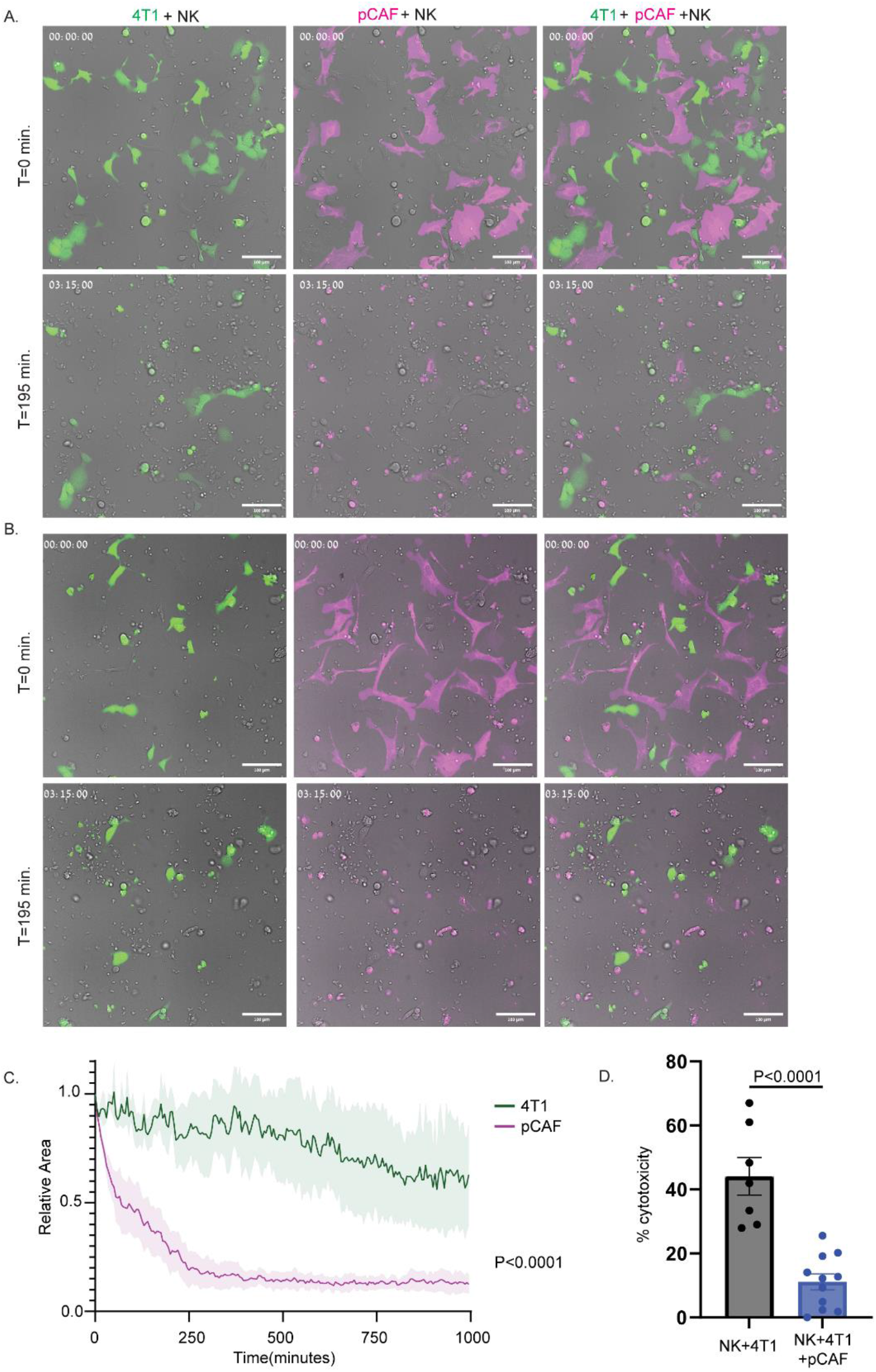
Live imaging of NK cell cytotoxicity in tri-cultures reveals enhanced susceptibility of CAFs to NK-mediated lysis. (A-B) Representative images of tri-cultures consisting of 4T1-GFP cells (green), pCAFs (magenta), and activated NK cells (unlabeled) at time point 0 and following approximately 3 hours of incubation from two independent experiments. pCAFs were freshly isolated from 4T1 tumors and NK cells were isolated from syngeneically matched BALB/c mice and were activated 24 hours prior to the experiment. (C) Quantification of 4T1 and pCAF death from live imaging experiments. Average cell area of either 4T1 or pCAF labeled and segmented cells was measured for each time point (see Methods). The average cell area for each time point was normalized relative to the average cell area at time 0. The graph summarizes data of 3 independent biological experiments, and displays the relative average cell area of 4T1 or pCAF cells for each time point, ±SEM. Statistical analysis was conducted using two-way ANOVA with time and cell type as discrete factors. The P value is provided for Time * Cell type interaction. (D) Quantification of cytotoxicity of co-cultures consisting of activated NK cells and 4T1-LUC cells or tri-cultures consisting of NK cells, 4T1-LUC cells, and isolated pCAFs from 4T1 tumors. 50k NK cells were activated for 24 hours with IL-2/15 and then seeded onto plates consisting of 10k 4T1-LUC cells or 10K 4T1-Luc cells and 10K pCAFs. Percent cytotoxicity is presented relative 4T1-LUC cells seeded alone. N=7 and N=11 biologically independent experiments for co cultures and tri cultures, respectively.

These data imply that CAFs can compete with cancer cells for interaction with NK cells via NKG2D/DNAM-1 receptor-ligand engagement, which subsequently hinders their ability to kill cancer cells.

### CAFs activate transcriptional programs regulating NK cell trafficking and adhesion

To investigate additional potential NK:fibroblast crosstalk mechanisms that may promote NK cell migration and sequestration into CAF rich areas *in-vivo*, we conducted bulk RNA-seq of tumor infiltrating NK cells (tiNK) *vs* splenic NK cells (sNK) from 4T1 tumor bearing mice. Differential expression analysis highlighted 517 genes upregulated and 237 genes downregulated in tiNKs *vs* sNK cells from spleens of tumor-bearing mice (Supplementary Table 1 and 2 and Supplementary Figure 9C-D). Pathway analysis revealed that tiNKs were characterized by regulatory activation and cell-death pathways and upregulation of checkpoint molecules such as *Ctla-4*^71^, *Lilr4b* ^72^, *Sult2b1*^73^ and *Cish*^74,75^. Genes involved in augmentation of adaptive immunity, such as *Ccl3* and *Ccl5,* were downregulated in tiNKs, whereas genes involved in dysfunctional stress modules, such as *Nr4a1* and multiple heat shock genes (*Hspa1a, Hspa1b, Hspa5, and Hspa13*^76^ were upregulated in tiNKs compared to sNK cells (supplementary Table 1). Moreover, *Rgs1* and *CD69*, recently shown to distinguish between circulating, tissue and tumor-infiltrating NK cells in different human cancers^76^ were significantly upregulated in tiNKs, whereas *Klf2* which marks circulating NK cells was increased in sNK cells.

Since we observed rapid NK migration to and cytotoxicity of CAFs in live imaging experiments, we hypothesized that fibroblasts may be rewired in cancer to promote NK cell migration and retention in stromal areas. To test this, we generated a curated list of activating, adhesion, and chemotactic ligands for NK cell receptors and assessed their expression levels on NMFs compared to CAFs in the 4T1 model (Supplementary Figure 9E). Both CAF subtypes demonstrated a global upregulation of the majority of these ligands compared to NMFs, with pCAFs exhibiting the highest expression of most NK ligands. These ligands may interact with chemokine receptors upregulated on tiNKs, such as Cxcr6, Cxcr4, Ccr7, and Ccrl2.

To assess the potential of CAFs in human tumors to compete with cancer cells for NK cell recruitment and retention, we renalyzed publicly available single-cell RNA-seq datasets of human breast cancer^77^ (Supplementary Figure 10A-B). We generated a list of ligands for human NK cell receptors and assessed their expression levels in CAFs compared to other cell types in the TME (Supplementary Figure 10). We found that fibroblast subsets consistently upregulated multiple chemotactic NK ligands compared to cancer cells (e.g. *RARRES2*, *CXCL12*, *CXCL1-3, CCL2-5*). In addition, ligands for NK adhesion molecules including *ICAM-1* and *VCAM-1* were more highly expressed in CAFs compared to cancer cells, and ligands for various NK activating receptors were predominantly expressed in CAFs (e.g. *NID1, CLEC2B*, and *CALR* which bind NKp444, NKp80, and NKp46, respectively^78–80^. These data collectively indicate upregulation of multiple genes by CAFs that may dictate NK migration to stromal areas, and underscore the relevance of our findings in murine models to human disease.

### Stromal expression of Nectin2 is associated with poor disease outcome in breast cancer patients

To determine the clinical significance of our findings, we stained for NK cells and their ligands in a cohort of 72 TNBC patients containing long-term clinical follow up^14^ (Figure 6). First, we sought to determine the prevalence of NK cells in stromal *vs*. tumor areas in these patients. To this end, we conducted immunohistochemical staining of NK cells combined with hematoxylin Eosin (H&E) in patient tumor microarrays (TMAs). We used image analysis to quantify the number of NK cells per tumor. We found relatively low amounts of NK cells in patient samples, in accordance with their known modest infiltration of solid tumors^81^. We then applied machine learning on the H&E staining to classify stroma-rich regions vs. cancer-rich regions, and used this classification to calculate the amount of NK cells per region. Across patients, NK cells were significantly enriched in stroma-rich regions vs. cancer-rich regions (Exact Wilcoxon rank sum test p−value = 0.00005, Figure 6A-B, Supplementary Table 3). This result may indicate that, similar to our findings in mouse models, NK cells interact with stromal cells also in human cancer.

**Figure 6.**
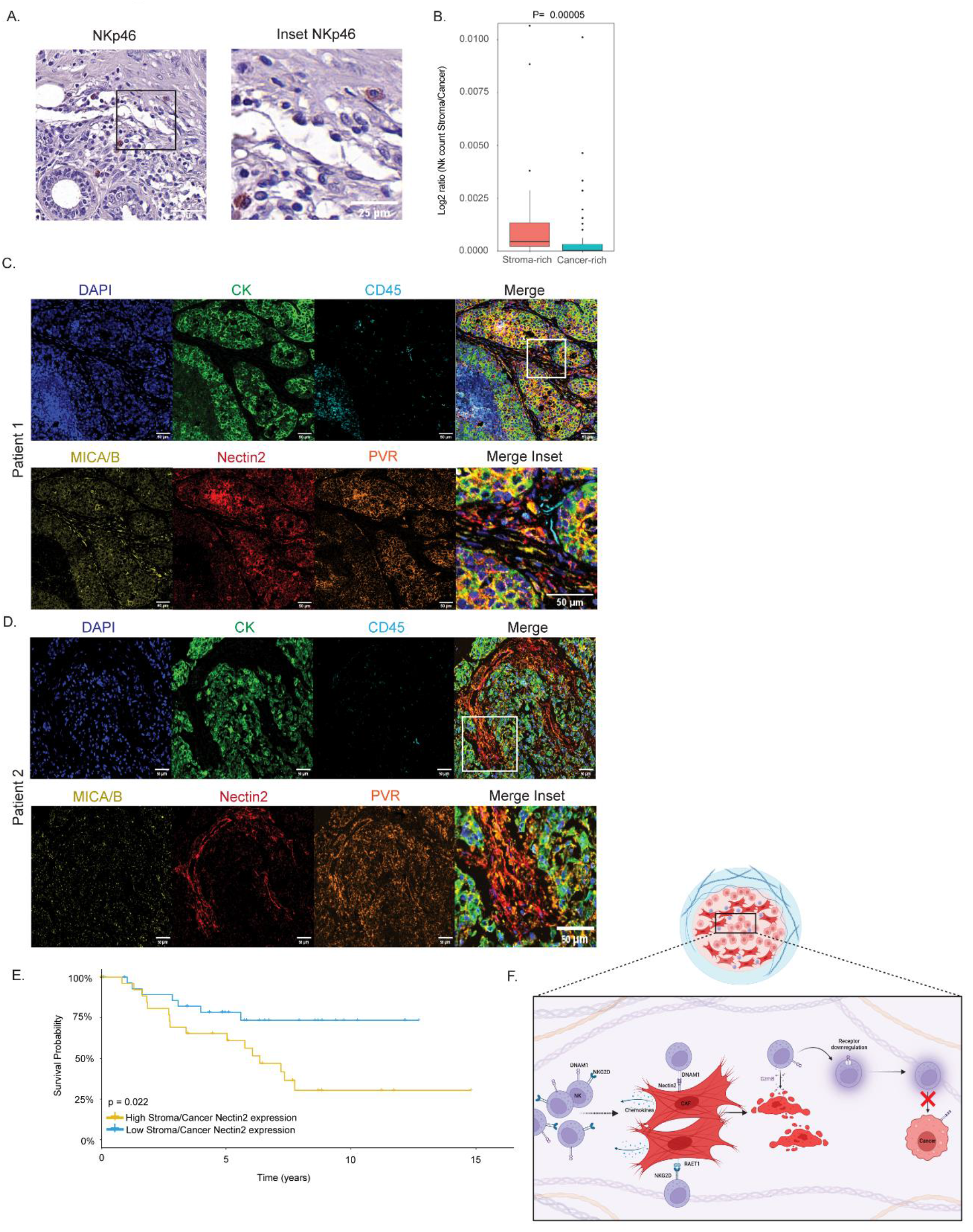
Stromal expression of the NK ligand Nectin2 is associated with poor disease outcome of breast cancer patients. (A) Representative IHC staining of NK cells (NKp46) in TMAs of TNBC patient cohorts. (B) Quantification of IHC images was conducted on the proportion of NK cells in stromal areas compared to cancer specified regions (Exact Wilcoxon rank sum test p−value = 0.00005) (C-D) Representative immunofluorescence images of 2 patient TMAs. Staining was conducted on DAPI, cytokeratin (cancer cells), CD45 (immune cells), and NK cell ligands NECTIN2, PVR, and MICA/B. (E) Kaplan-Meir survival analysis of patients stained in (C-D). Analysis was conducted on the ratio of CAFs expressing NECTIN2 out of the total CAFs, compared to the ratio of positive cancer cells out of total cancer cells, with groups stratified above (high) or below (low) the median value. Statistical analysis was conducted using two-sided log-rank test. (F) Scheme of proposed model. Reduction of NK cell receptors following engagement with CAFs may result in impaired cytolysis of cancer cells by NK cells in the TME

We therefore next assessed the expression of NK-activating ligands on CAFs in the same patient cohort. We stained the TMAs with antibodies for the NK ligands Nectin2, PVR, and MICA/B, and with antibodies for cancer cells (CK) and immune cells (CD45) Figure 6C-D, Supplementary table 3). All three ligands were expressed on CK^+^ cancer cells as well as on CK^-^CD45^-^ stromal cells (Figure 6C), and the expression of PVR and NECTIN2 was more abundant in cancer than in stroma, while the expression of MICA/B was similar between both groups (Supplementary Figure 11A). To test the potential clinical significance, we conducted Kaplan Meier survival analysis. Stratification of patients based on expression of the ligands in the total tumor area did not reveal significant correlation with disease outcome (Supplementary Figure 11B-D). When stratified by the ratio of ligand expression in stroma *vs.* tumor, however, we found that a high ratio of stroma/tumor expression of Nectin2 was associated with significantly poorer overall survival (P=0.022, Figure 6E). Collectively, these data emphasize the importance of spatial context of ligand expression in the TME. Furthermore, they support the hypothesis that CAFs in human breast cancer may interact with NK cells through Nectin2::DNAM-1, and that these interactions could lead to inhibition of NK cell anticancer activity resulting in poor disease outcome.

## Discussion

Accumulating evidence has emerged in recent years regarding the role of CAFs in subversion of immune responses against cancer^4^, yet the regulation of NK cells by CAFs has remained relatively unexplored. Here, using mouse models of breast cancer, we find a mechanism through which CAFs can impede NK cell cytotoxicity against cancer cells, by acting as ‘decoys’ for NK cell killing. Specifically, we demonstrate that CAFs can upregulate ligands for two critical NK cell activating receptors: NKG2D and DNAM-1. In the presence of CAFs, the surface expression of NKG2D and DNAM-1 on NK cells is dramatically reduced, and the ability of NK cells to mount cytotoxic responses against cancer cells is inhibited (Figure 6F). We further show that NK cells are enriched in stromal areas of human tumors and that expression of their ligands in human breast cancer stroma is associated with poor survival, highlighting the potential clinical relevance of our findings.

One mechanism impeding the capability of NK cells to mount successful responses against cancer involves downregulation of activating receptors, including NKG2D and DNAM-1 in the TME^82^. Here we propose that not only the cancer cells but also the CAFs express ligands for the NK activating receptors and may be dominant players in mediating inhibition of NK cell cytotoxicity. These results are in accordance with previous observations, showing that engagement of activating receptors with their cognate ligands expressed on cancer cells leads to internalization and degradation of the receptors and inhibition of cytotoxicity^57–63,83^. Specifically, we show that Nectin2 is expressed on human CAFs and this expression is correlated with poor disease outcome. In mice we find that CAFs can upregulate Nectin2, PVR, RAE-1, H60a, and MULT1, leading to reduction in surface expression of NKG2D and DNAM-1 and resulting in loss of cytotoxicity.

Other mechanisms mediating CAF-NK cell interactions may very well be in play. These could include secretion of soluble NKG2D and DNAM-1 ligands^84,85^ as well as interactions between other NK receptors and their ligands. One example may be NKp46, which we observed to be partially downregulated in the presence of CAFs. Recently, NKp46 was demonstrated to promote NK recognition of cancer cells via externalized calreticulin, exported from the ER during ER stress^80^. We and others have previously shown that stress responses induced in the TME are conferred to CAFs, with critical ramifications for cancer progression^16, 48,86,87^. It is plausible that CAFs express or secrete ligands for NKp46 leading to its downregulation. An additional mechanism that may mediate CAF regulation of NK cells may operate through chemotactic and adhesion molecules. Indeed, RNA-seq data from the 4T1 mouse model and from cancer patients revealed upregulation of chemotactic ligands that may sequester NK cells in CAF-rich areas within the TME. Furthermore, histological staining of NK cells from patient samples revealed a significantly higher enrichment of NK cells in stromal areas.

NK cell interactions with activated fibroblasts were observed under different pathological conditions. During liver fibrosis, for example, activated hepatic cells are targeted by NK cells via NKG2D^67^. NK cells may also play a role in targeting Fibroblast-like synoviocytes during Rheumatoid arthritis, and thus play a role in local inflammation^68^. In context of cancer, hepatic stellate cells contribute to breast cancer metastases formation by inhibiting NK cells through secretion of CXCL12^88^; these data strengthen our findings in scRNA-seq human TNBC data showing that CAFs are the major source of CXCL12 in the breast cancer TME (Supplementary Fig.10G). Activated NK cells were also reported to target and lyse pancreatic stellate cells in addition to pancreatic cancer cell lines^89^. Under physiological wound healing, upregulation of NKG2D and DNAM-1 ligands by activated fibroblasts may act as a timer for regulating fibrotic processes. This is concordant with the observation that NMFs demonstrate a striking increase in these ligands following culture ex-vivo, which induces their activation and transition to a myofibroblastic state^64–66^. In cancer, it may very well be that NK cell involvement in the resolution of fibrosis is rewired and manipulated by the TME. Specifically, fibrotic signatures induced in CAFs may promote a chronic stimulatory state in NK cells, and possibly CD8+ T-cells, to expedite an exhausted phenotype.

Multiple therapies for solid tumors are currently being developed and some clinically tested which target NKG2D and DNAM-1 signaling, including chimeric antigen receptor (CAR)-expressing cells and multiple antibody constructs^90,91^ (NCT05213195, NCT05776355, NCT05378425) that are aimed towards cancer cells with high expression of NKG2D and DNAM-1 ligands. The results shown here may warrant screening patients for ligand expression patterns in stromal cells in tumors prior to treatment, since it is conceivable that high stromal expression of these ligands may attract effector cells to stromal areas instead of the cancer areas. Indeed, we find that high expression of Nectin2 in stromal cells compared to cancer cells is associated with poor disease outcomes in patients. In a previous study we also observed a sharp increase in stromal expression of Nectin2 in BRCA2 mutated pancreatic cancer patients, with inhibitory effects on cytotoxic CD8+ T-cells^16^, raising the possibility of its use as a stromal biomarker in different cancers. Thus, the spatial organization and quantity of CAFs in tumors may dictate reduction of cytotoxic lymphocyte activity via the NKG2D-DNAM-1 axis, since both NK cells and CD8+ T-cells rely on these receptors for immune-surveillance^92,93^. On the other hand, the observation that CAFs upregulate NKG2D and DNAM-1 ligands also opens the possibility of utilizing activated NK cells to modulate solid tumors that are characterized by high desmoplasia as a combinatorial immunotherapeutic approach; One interesting avenue may incorporate adoptive NK cell administration followed by conventional checkpoint blockade that may initially reduce stromal content, and therefore provide potent synergistic effects in solid tumors. We hope that our findings here may provide additional understanding of the complex immune regulation by CAFs in tumors, and enable the development of new approaches to target immune-CAF circuits in the TME.

## Methods

### Ethics Statement

All clinical data were collected following approval by the Sheba Medical Center Institutional Review Board (IRB), protocol no. 8736-11-SMC. All animal studies were conducted in accordance with the regulations formulated by the Institutional Animal Care and Use Committee of the Weizmann Institute of Science (protocols 02820323-2, 04140522-1, 02590321-1, 08801120-2).

### Human patient samples

TMAs from 72 TNBC patients (three cores per patient), with matching H&Es were retrieved from the archives of Sheba Medical Center under IRB no. 8736-11-SMC. All clinical data were collected following appropriate ethical approvals. Approval was given by the Sheba Medical Center IRB (protocol no. 8736-11-SMC) with full exemption for consent form for anonymized samples.

### Mice

BALB/c and C57BL/6 mice were purchased from Envigo and maintained at the Weizmann Institute’s animal facility under specific pathogen-free conditions. Mice were sacrificed by cervical dislocation for tumor or normal tissue harvesting.

### Cancer cell lines

4T1 cells expressing firefly luciferase (pLVX-Luc) were kindly provided by Zvika Granot’s lab (HUJI). E0771 cells were kindly provided by Ronen Alon’s lab (WIS). GFP-expressing 4T1 cells (4T1-GFP) were generated using the FUW-GFP vector and mCherry-luc-expressing E0771 cells were generated using a luc2a-mcherry vector. 4T1 and E0771 cells were cultured in Dulbecco’s modified Eagle’s medium (DMEM; Biological Industries, 01-052-1A) with 10% fetal bovine serum (FBS; Invitrogen).

### Primary NK Cell isolation and culture

Primary murine NK cells (pNKS) were isolated from spleens of BALB/c and C57BL/6 mice using magnetic separation (Miltenyi mouse NK Cell Isolation Kit). NK cells were validated for purity utilizing FACS positive staining for CD45 (immune cells), negative staining for CD3 (T-cells), and positive staining for NKp46 and CD49b (NK cells). NK cells were maintained in RPMI 1640 (Biological Industries, 01-100-1 A) supplemented with 0% FBS, 1% MEM NEAA, 1% 0.5 M HEPES buffer, 1% L-glutamine, 1% sodium pyruvate and 0.0004% βM-EtOH (Biological Industries). For activation, NK cells were supplemented with 1000 u/ml of recombinant human IL-2 and 100 ng/ml of recombinant human IL-15 for 24 hours.

### Orthotopic injection to the mammary fat pad

8 Week old BALB/c or C57BL/6 female mice were injected under anesthesia with 100,000 4T1-GFP cells or 600,000 E0771-mCherry cells reconstituted in 50 μl of PBS, into the lower left mammary fat pad.

### Normal mammary fat pad isolation and dissociation

NMFs were collected from healthy 8 week old BALB/c females as previously described^2,14^. Fat pad tissue was minced and dissociated using the gentleMACS dissociator with an enzymatic digestion solution consisting of 1 mg ml−1 collagenase II (Merck Millipore, 234155), 1 mg ml−1 collagenase IV (Merck Millipore, C4-22) and 70 U ml−1 DNase (Invitrogen, 18047019) in DMEM. The samples were filtered through a 70-μm cell strainer into ice-cold MACS buffer (PBS with 0.5% BSA) and cells were pelleted by centrifugation at 350g for 5 min at 4 °C.

### Primary CAF isolation

Primary CAF isolation was conducted as previously described^2,14^. Briefly, 6 weeks following 4T1-GFP or E0771-mcherry injection, animals were euthanized, and tumors were excised and dissociated, and incubated with enzymatic digestion solution consisting of 3 mg ml−1 collagenase A (Sigma Aldrich, 11088793001) and 70 U ml−1 DNase in DMEM at 37 °C using the standard gentleMACS solid tumor program. To enrich for stromal cells, single-cell suspensions were incubated with anti-EpCAM (Miltenyi, 130-105-958) and anti-CD45 (Miltenyi, 130-052-301) magnetic beads for elimination of residual immune and cancer cells, and transferred to LS columns (Miltenyi, 130-042-401). The stromal enriched (CD45, EpCAM depleted) flow-through was collected and pelleted for further FACS sorting.

### Flow cytometry and sorting of CAFs

CAFs were incubated in FACS buffer for 30 minutes at 4°C following staining with antibodies detailed in Supplementary Table 3. Following antibody incubation, cells were washed with a FACS buffer. Propidium iodide (PI) was added before FACS sorting to distinguish between live and dead cells. Gating strategies for pCAFs and sCAFs are shown in Supplementary figures 2A and 3A for 4T1 and E0771 models, respectively. The cells were sorted using a FACSAria Fusion sorter (BD Biosciences) into FACS tubes containing 1 mL of complete DMEM.

### Flow cytometry and sorting of NK cells

To sort for splenic and tumor infiltrating NK cells for RNA sequencing, NK cells from spleens and 4T1 tumors were incubated in FACS buffer for 30 minutes at 4°C following staining with antibodies detailed in Supplementary Table 4. Following antibody incubation, cells were washed with FACS buffer. Propidium iodide (PI) was added before FACS sorting to distinguish between live and dead cells. The gating strategy for NK cells is shown in Supplementary figure 1A. The cells were sorted using a FACSAria Fusion sorter (BD Biosciences) into FACS tubes containing 1 mL of complete RPMI.

### Bulk RNA-seq

RNA from sorted NK cells and fibroblasts was purified using the Dynabeads mRNA DIRECT Purification Kit (Thermo Fisher Scientific; cat. #61012). A total of 1 × 10^4^ pNKs or fibroblasts were isolated and sorted into 40 μL of lysis/binding buffer. RNA was isolated using the commercial protocol. Libraries were generated with the MARS-seq protocol^94^.

### Differential gene expression analysis

For the identification of differentially expressed genes between spleen-derived and tumor-infiltrating NK cells and between all fibroblast subsets, the R package DESeq2 pipeline was used^95^. Biological replicates were included in the design formula to adjust for batch effects. An adjusted P value <0.05 and an absolute log2-fold change >1 were used for statistical significance. Heat maps were generated using Morpheus, https://software.broadinstitute.org/morpheus.

### GO enrichment analysis

Gene-set enrichment analysis was performed using Metascape (http://metascape.org).

### NK-CAF co culture for assessment of NK cell receptor expression and cytotoxicity

Following isolation and FACS sorting, 70*10^4^ NMFs and CAFs were allowed to recover in RPMI for 4 days. RPMI was subsequently aspirated, and 0.5*10^6^ NK cells were resuspended in RPMI containing IL-2+IL-15 were co-incubated with the fibroblasts for 24 hours. Following co-incubation, plates were gently tapped and non-adherent NK cells were collected for FACS analysis of surface receptors and cytotoxicity assays against 4T1 or E0771 cancer cells. Unactivated and activated NK cells cultured alone served as negative and positive controls, respectively.

### Flow Cytometry of NK cell receptors and ligand expression on fibroblasts and TNBC cells

Cells subjected to flow cytometric analysis included NK cells cultured in the presence of CAFs, primary fibroblasts from tumors and naive fat-pad tissue, fibroblasts recovered in culture for 4 days, and TNBC cells (4T1 and E0771). Gating strategies for FACS analysis are presented in Supplementary Figure 1 A, 2A, and 3A. Cells were collected and stained with antibodies for activating or inhibitory cell surface receptors (in the case of NK cells) and NKG2D/DNAM-1 ligands (in the case of fibroblasts, 4T1, and E0771 cells) detailed in Supplementary Table 4. Staining was performed in ice cold FACS buffer for 30 min. Following incubation, cells were washed and receptor/ligand expression was analyzed via FACS analysis. Dead cells were excluded using Ghost Dye staining (TONBO biosciences, 13-0863-T500). FACS analysis was performed using flowjo software v.10.7.1. Analysis conducted included the geometric mean fluorescence intensity of all the live gated cell populations of interest.

### Cytotoxicity Assays

NK cells under all different treatment conditions were cultured at a 5:1 effector:target ratio (50K NK cell with 10K cancer cells or CAFs) for 6 hours at 37° Celsius and 5% CO2 in 96 flat well plates. Wells were subsequently lightly agitated and washed twice to remove NK cells. Next, wells were reconstituted with 50 microliters of RPMI medium. In the case of 4T1-Luc and E0771-Luc cells, cells were mixed with 50 microliters of Stead-Glo® (Promega, E2510) Luciferase Assay System buffer. In the case of fibroblasts, cells were mixed with 50 microliters of CellTiter-Glo® Luminescent Cell Viability buffer. Each experimental condition contained between 3-5 internal technical replicates that were averaged. Specific cytotoxicity % was calculated by normalizing all signals (sample well) to the cancer cells or fibroblasts cultured alone (without NK cells-control well) basal luminescence, utilizing the following formula:

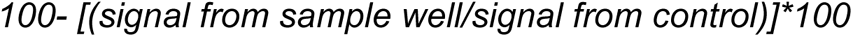

Negative values (indicating negative cytotoxicity) were replaced with a value of 0. In the case of 4T1 and E0771 cytotoxicity experiments, specific cytotoxicity was normalized to samples of NK cells activated alone (without CAFs). In the case of CAF cytotoxicity experiments, specific cytotoxicity was normalized to NK cells activated in the presence of IgG control.

### Conditioned media

70*10^4^ CAFs and NMFs were allowed to rest in RPMI following initial isolation for 24 hours. Subsequently, the medium was aspirated and replaced, and cells were allowed to grow for an additional 4 days. The medium was then collected and frozen at −80c for use on NK cells. Prior to incubation of NK cells with CM, CM was diluted at a 1:1 ratio with regular RPMI medium.

### NK cell receptor blockade

For NK cell receptor blockade experiments, NKG2D and DNAM-1 surface receptors on NK cells were blocked utilizing 25 μg/ml of purified anti-mouse NKG2D (Biolegend, clone CX5, 130202), purified anti-mouse DNAM-1 (Biolegend, clone 10E5, 128822), or Mouse IgG1 Isotype Control control (R&D, MAB002) for 15 minutes at room temperature prior to culture with cancer cells or CAFs. Following 15 minute incubation, NK cells were cultured with cancer cells or CAFs in the presence of each blocking antibody.

### CAF ligand blockade

For CAF ligand blockade experiments, CD155, Nectin2, and Pan-Rae-1 expressed on CAFs were blocked utilizing 25 μg/ml of anti-mouse pan Rae-1 (R&D,AF1136), Ultra-LEAF™ anti-mouse CD155 (Biolegend, 942103), anti-mouse Nectin2 (R&D, MAB3869), or Mouse IgG1 Isotype Control control (R&D, MAB002) for 15 minutes at room temperature prior to culture with NK cells. Following 15 minute incubation, NK cells were cultured with CAFs in the presence of each blocking antibody.

### Live imaging experiments and analysis

10K 4T1-GFP cells and 10K pCAFs stained with Cell Tracking Dye Kit - Red - Cytopainter (ab138893) were co-cultured at a 1:1 ratio for tri-culture experiments with NK cells. For co culture experiments (4T1 or pCAF cells alone with NK cells) 20K 4T1-GFP or stained pCAFs were seeded. NK cells from spleens of BALB/C mice were activated overnight with IL-2+IL-15, and 50K NK cells were introduced to wells with 4T1/pCAFs and immediately imaged for 16 hours.

Live cell images were acquired every 5 min using a spinning disk confocal Eclipse TI2 microscope with CSU-W1 50µm pinhole (Nikon). Images were acquired with CFI Plan Fluor ×20/0.75 objective, GFP and Cy5 cubes and with a prime BSI Scientific CMOS camera (Teledyne Photometrics).

Images were segmented using Cellpose 2.0^96^ with a cyto2 pretrained model for the GFP channel and a cyto2 retrained model for the magenta channel. Areas of the segmentations were calculated using Fiji image processing platform^97^.

The Area for each segmented cell in the GFP or magenta channel was calculated using the Fiji TrackMate plugin for each of the 200 frames. In each frame, the average area of all cells was calculated. Each averaged area in each frame for the GFP and magenta channels was normalized to the area of the first frame to generate temporal graphs of relative cell death.

### Single cell analysis

A publicly available human Breast cancer single-cell dataset (GSE161529) was analyzed using the Seurat (V4.0) R toolkit. BRCA1 and male samples were filtered out. In addition to the Seurat pipeline, Harmony integration^98^ with default parameters was used to minimize the patient batch effect, and shared nearest neighbor (SNN) modularity optimization-based clustering was then used with a resolution parameter of 0.07. A COL1A1/DCN positive cluster (CAFs and Normal fibroblasts - 24942 cells) was pulled and reanalyzed with the same pipeline (clustering resolution of 0.2). After labeling the CAF subclusters, the labels were projected on the original cells in the full data UMAP.

### Immunohistochemistry of NK cells in human tissues

4 μm FFPE sections from the human TNBC TMA were deparaffinized and treated with 1% H_2_O_2_. Antigen retrieval was performed using Tris-EDTA. Slides were blocked with 10% normal horse serum (Vector Labs, S-2000) and the NK antibody NKp46 (listed in Supplementary Table 4) was used. Visualization was achieved with 3,3’-diaminobenzidine as a chromogen (Vector Labs, SK4100). Counterstaining was performed with Mayer hematoxylin (Sigma Aldrich, MHS16). TMA slides were scanned by the Pannoramic SCAN II scanner, with 20×/0.8 objective (3DHISTECH, Budapest, Hungary).

### Immunofluorescent staining of human tissues

FFPE sections from the human TNBC TMA were deparaffinized and incubated in 10% neutral buffered formalin (prepared by 1:25 dilution of 37% formaldehyde solution in PBS). Antigen retrieval was performed with citrate buffer (pH 6.0; CD45 and CK) or with Tris-EDTA buffer (pH 9.0; PVR, NECTIN2, MIC A\B and PDPN). Slides were blocked with 10% BSA + 0.05% Tween 20 and the antibodies listed in Supplementary Table 4 were diluted in 2% BSA in 0.05% PBST and used in a multiplexed manner using the OPAL reagents (Akoya Biosciences). Briefly, following primary antibody incubation, each one overnight at 4 °C, slides were washed with 0.05% PBST, incubated with secondary antibodies conjugated to HRP, washed again and incubated with OPAL reagents. Slides were then washed and antigen retrieval was performed as described above. Then, slides were washed with PBS and stained with the next primary antibody or with DAPI at the end of the cycle. Finally, slides were mounted using Immu-mount (#9990402, Thermo Scientific). We used the following staining sequences: CD45 → CK → Nectin2 → PVR → MIC A\B → PDPN → DAPI. Each antibody was validated and optimized separately and then MxIF was optimized. TMA slides were imaged with PhenoImager®-Fusion 2.0 upright scanner.

### Image analysis - MXIF

First, spectral unmixing and background subtraction were performed by creating an algorithm for this specific staining and tissue (tumor, breast) using inForm (version 2.8; Akoya Bioscience). Next, TMA cores were stitched using QuPath^99^ (version 0.4.0). Quantification of TMA staining was performed using QuPath (version 0.3.4). TMA cores were defined using QuPath. Cores that were left with less than 1/3 of the tissue due to damage or disruption in the staining process were excluded from the analysis. This exclusion criterion also applied to cores containing mostly regions of adipose tissue due to nonspecific staining. Object-based analysis was then performed. Cells were segmented based on nuclear staining (DAPI) using cellpose^100^. Next, we trained the ‘Random Trees’ classifier to categorize cells for different markers or to ignore them based on the specific channel of interest, independently of all the other markers, in addition to morphology features, DAPI and autofluorescence channels. After training a classifier for each marker, a composite classifier was created and applied to all images. Positive cells for a marker represent the percentage of positive cells in each core relative to the total cell count in each core. Cancer cells were defined by CK^+^ classifier and stromal cells were defined as CK^-^CD45^-^. PDPN staining was also performed to assess pCAFs, which are a subset of the CAFs in breast cancer, but eventually we did not proceed with the PDPN^+^ analysis to assess the effects of the entire CAF population on patient outcomes. The patient score was called through the averaging of 2-4 cores. To test for the correlation with survival we used a Kaplan Meier analysis with a log rank test.

### Image analysis - IHC

IHC stained slides were scanned using the Pannoramic SCAN II scanner with a 20×/0.8 objective (3DHISTECH, Budapest, Hungary). The quantification of NK^+^ cells and their distribution (stroma/cancer) within the tumor area of each core was performed using QuPath (version 0.4.0) through pixel classification and object detection. Briefly, a pixel classifier was applied to identify tissue areas, with damaged areas (folding, tearing) and adipose tissue classified as ‘ignore.’ Another pixel classifier was applied to identify cancer-rich areas marked as ‘cancer’ and stroma-rich areas as ‘stroma.’ Subsequently, cells were defined through nuclei segmentation, and the expansion of the nuclei was assumed for cell identification. The StarDist method, employing the ‘he_heavy_augment’ model, was used for nuclei segmentation. The classifier method utilized was the Artificial Neural Network (ANN_MLP) with moderate resolution, and the same classification parameters were applied to all images. The number of NK^+^ cells in the stroma or cancer areas was quantified based on the total cell count per core^101^. Cores with less than 1/3 of the tissue remaining due to damage or disruption in the staining process were excluded from the analysis. This exclusion criterion also applied to cores predominantly containing regions of adipose tissue due to nonspecific staining. To test for differences in the ratio of cells within stroma and tumor we applied an exact Wilcoxon test.

### Statistics and reproducibility

Statistical analysis and visualization were conducted utilizing R (Versions 3.6.0 and 4.0.0, R Foundation for Statistical Computing Vienna, Austria) and Prism 9.1.1 (Graphpad, USA). Student’s t-test (comparing two groups) or ANOVA (comparing more than two groups) were used to analyze normally distributed data. Statistical tests were defined as significant using a p value < 0.05 or an FDR < 0.05 for multiple comparisons. “ns” in all Figures indicates p-values greater than 0.05.

## Data availability

RNA sequencing data of murine NK cells and sCAFs were deposited in Gene Expression Omnibus (GEO) and can be accessed under GSE247218. Murine RNA sequencing data of NMFs and pCAFs were previously reported by us^2,30^ and available via GSE195865. Single cell RNA-seq of human patient data^77^ is available under (GSE161529). All additional data supporting the findings of this study are available upon reasonable request.

