## Supplementary Figures for "Cancer-associated fibroblasts serve as decoys to suppress NK cell anti-cancer cytotoxicity"

Ben-Shmuel et al. Supplementary Figure 1

A.

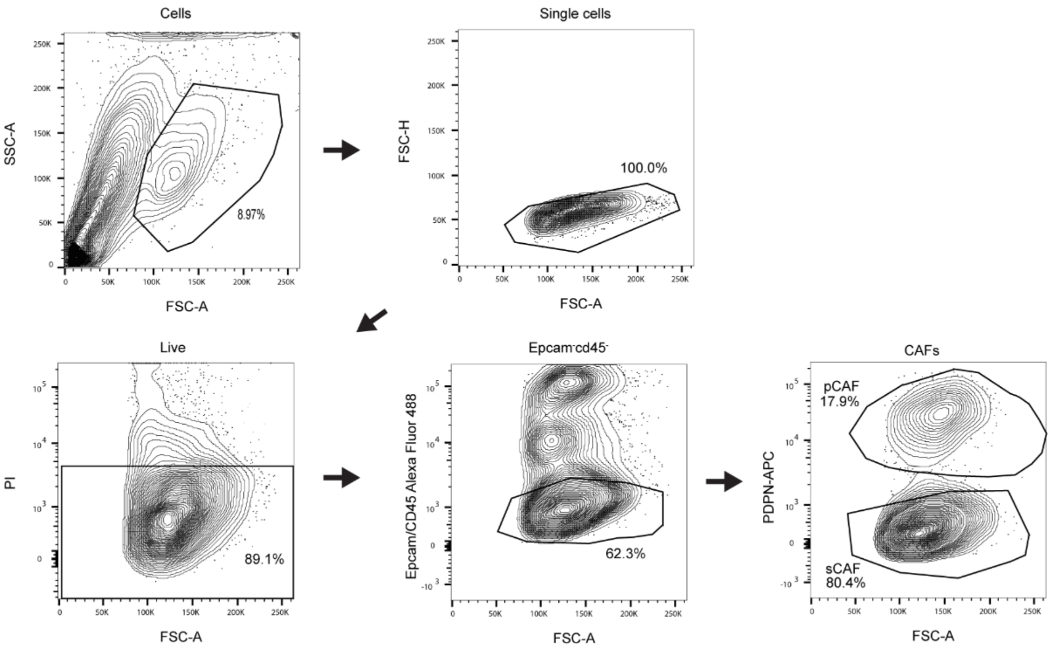


B.

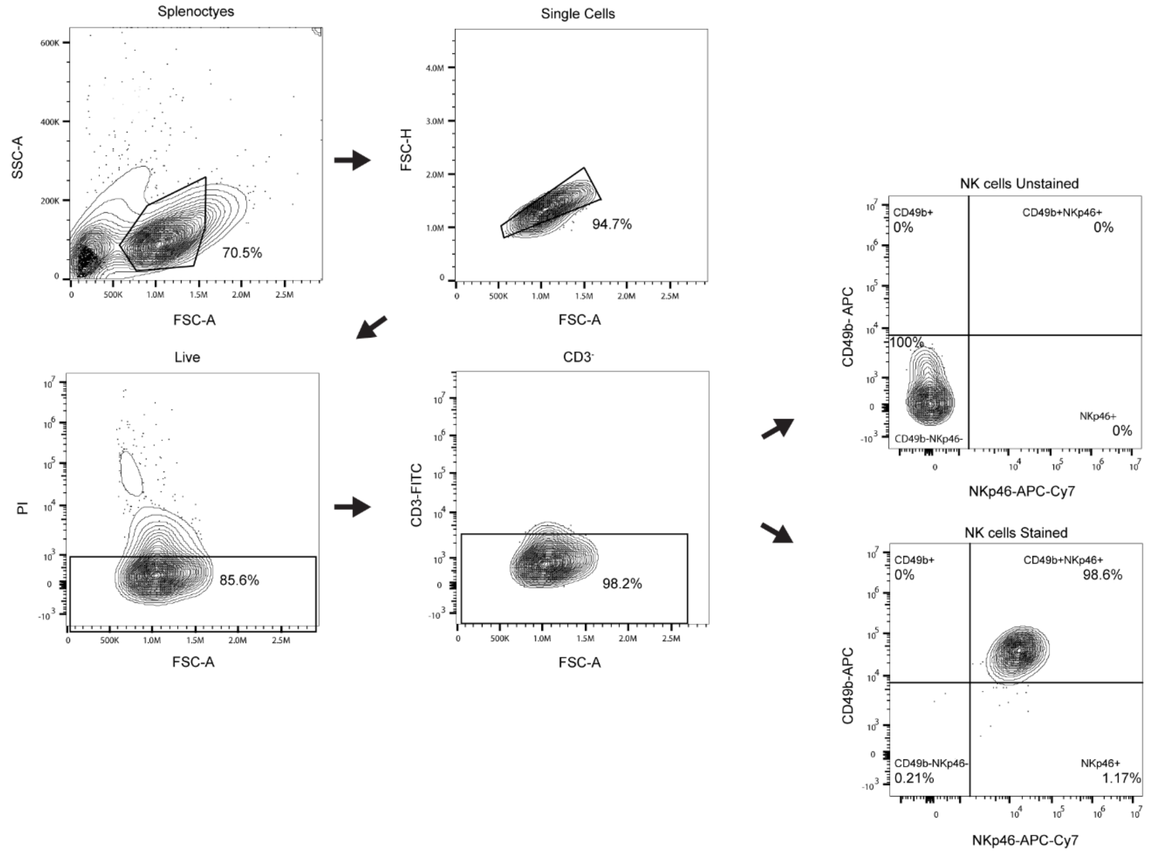

**Supplementary Figure 1. Gating strategy for primary CAFs and pNKs from BABL/C mice.** A. 4T1-GFP tumors were dissociated as described in the methods section. CAFs were selected for based on negative PI live/dead staining and negative staining for epithelial and immune cell markers (GFP marking cancer cells, EPCAM, and CD45). pCAFs and sCAFs were isolated as previously described, based on positive or negative staining for podoplanin (PDPN), respectively. B. NK cells were isolated from spleens of BALB/C mice using magnetic separation (see methods). Enrichment of NK cells was assessed by negative staining for CD3 (T-cells) and positive staining for Nkp46 and Cd49b.

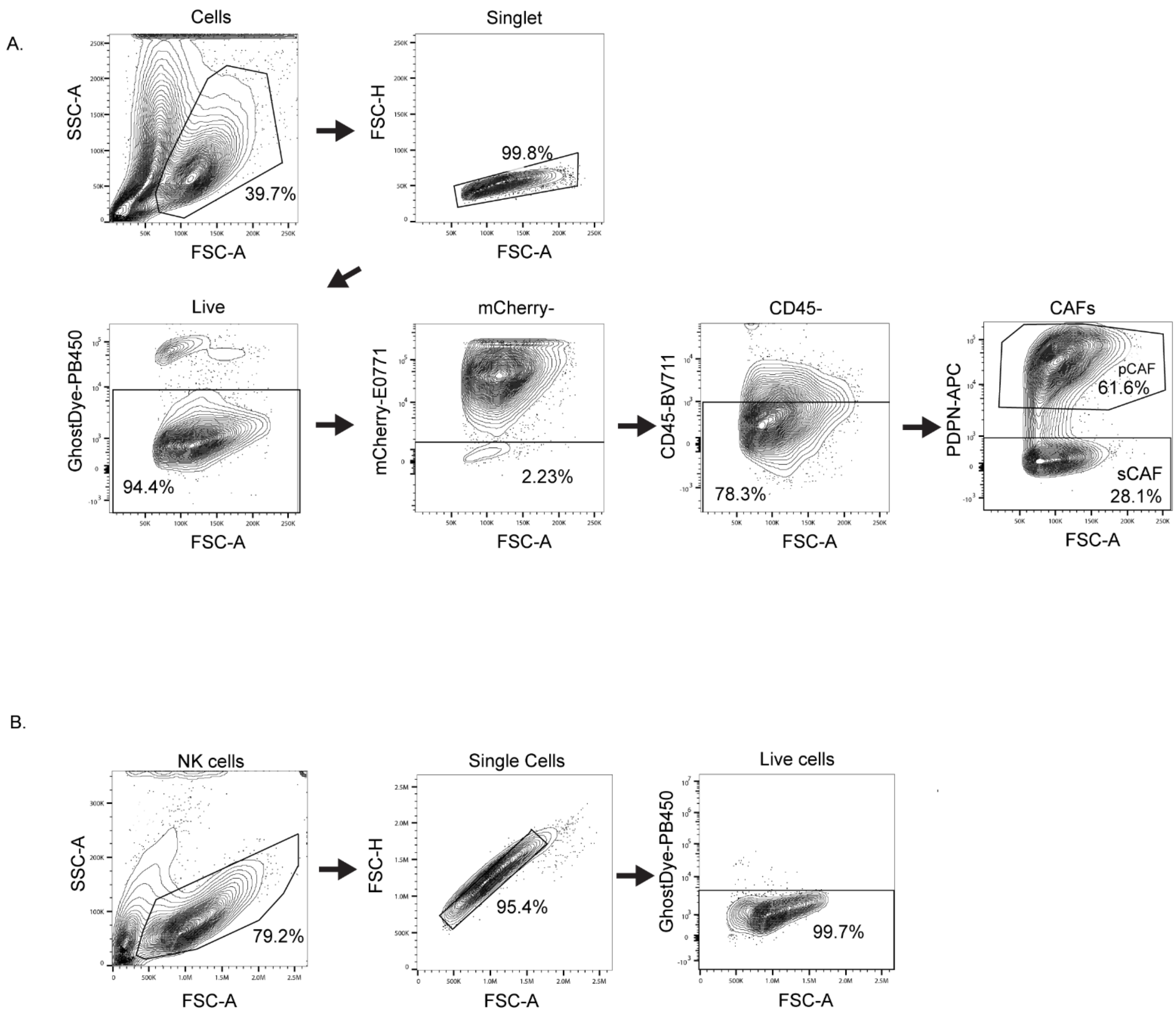


**Supplementary Figure 2. Gating strategy for primary CAFs from E0771 tumors and NK cells for FACS analysis.** A. E0771 tumors were dissociated as described in the methods section. CAFs were selected for based on negative GhostDye live/dead staining and negative staining for epithelial and immune cell markers (mCherry marking cancer cells and CD45). pCAFs and sCAFs were isolated based on positive or negative staining for podoplanin (PDPN), respectively. B. Gating of NK cells following culture with CAFs for assessment of surface receptor expression.

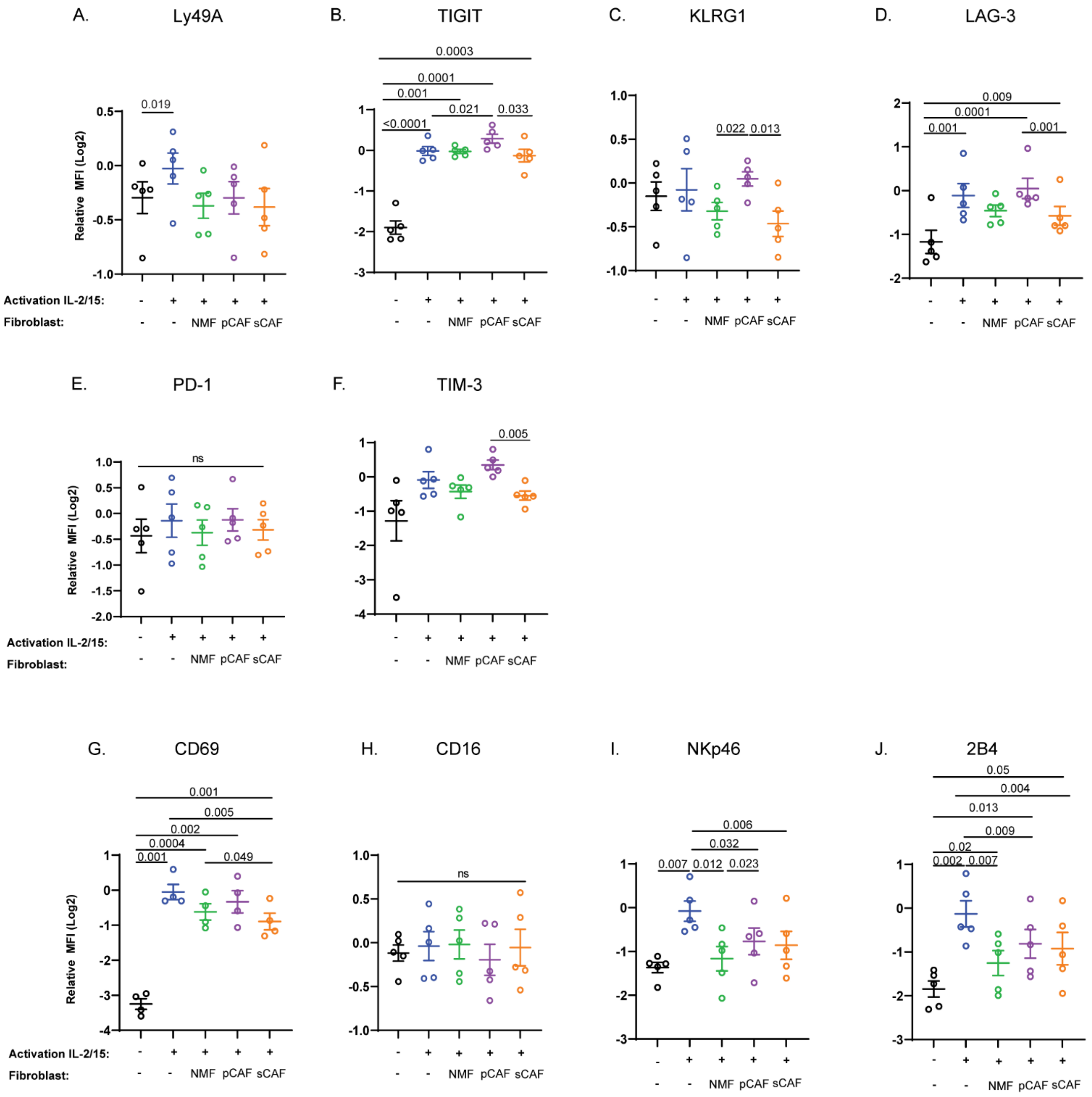


**Supplementary Figure 3. Inhibitory NK cell receptor expression does not change following CAF coculture.** A-J. Quantification of NK receptor surface expression. MFI (Log2 transformed) of each receptor is shown for all live gated NK cells. Data is presented as mean ± SEM of N=5 independent biological experiments relative to NK cells activated alone.

Ben-Shmuel et al. Supplementary Figure 4

A.

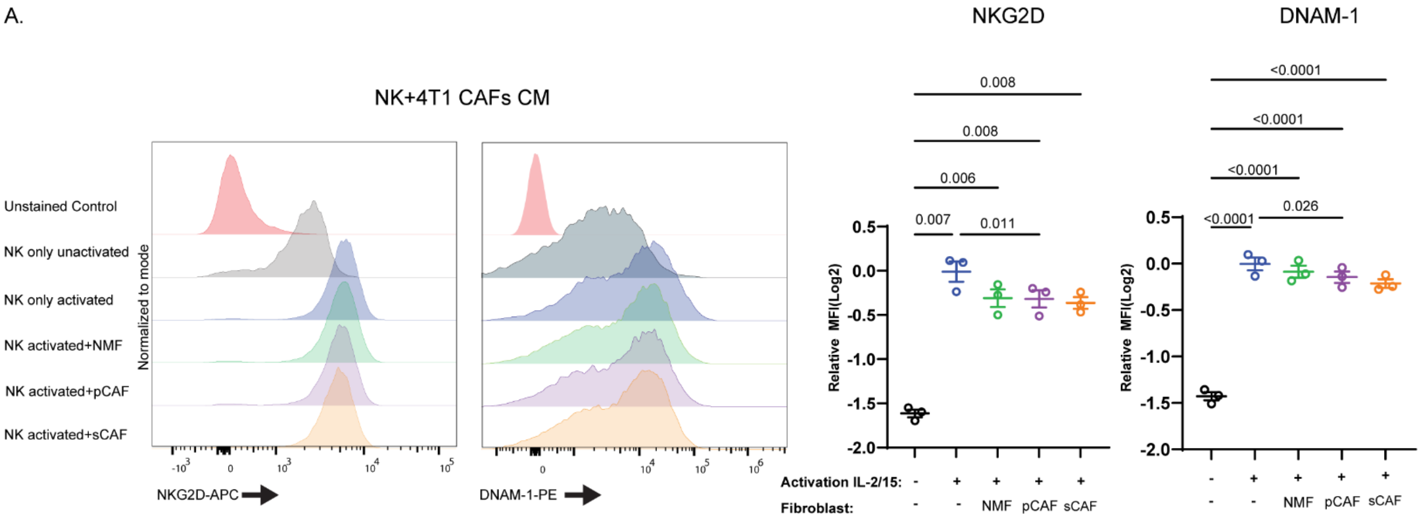


B.

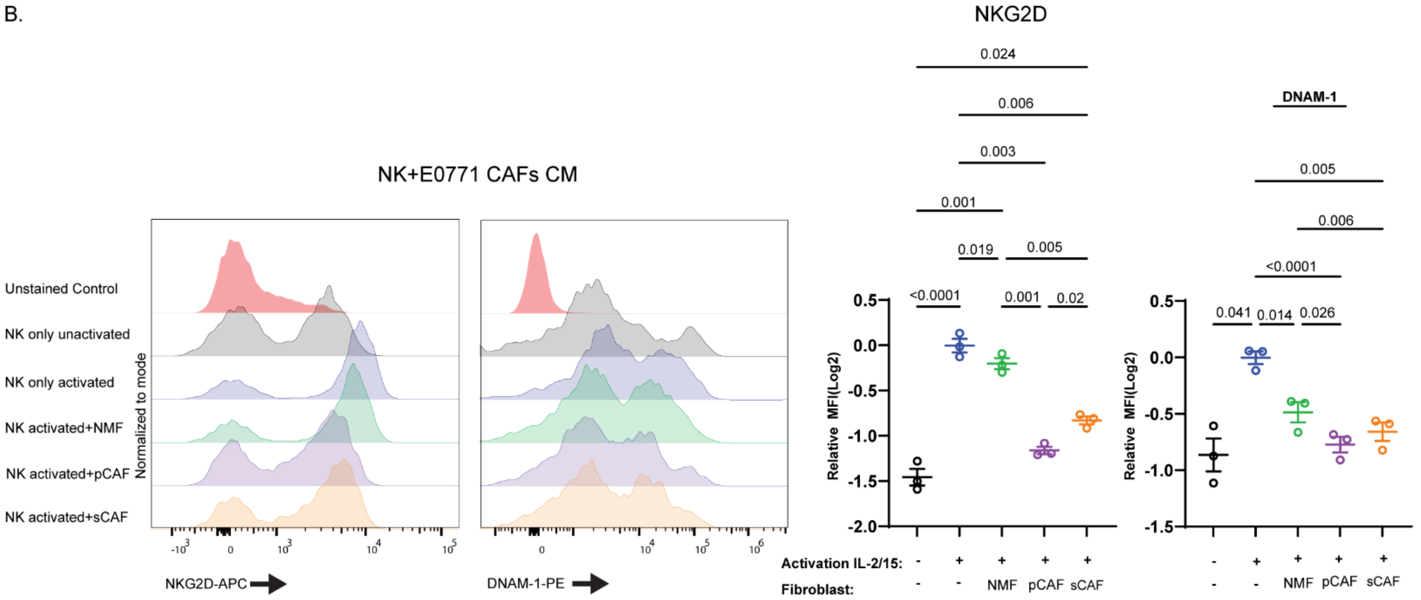

**Supplementary Figure 4. 4T1 CAF CM does not affect NK cell receptor expression** A. Representative FACS plot and quantification of NKG2D and DNAM-1 expression on NK cells from BALB/C mice following activation in regular medium or CM from NMFs, pCAFs, or SCAFs isolated from 4T1 tumors. B. Representative FACS plot and quantification of NKG2D and DNAM-1 expression on NK cells from C57BL/6 mice following activation in regular medium or CM from NMFs, pCAFs, or SCAFs isolated from E0771 tumors. MFI (Log2 transformed) of each receptor is shown for all live gated NK cells. Data is presented as mean ± SEM of N=3 independent biological experiments relative to NK cells activated with control medium. FACS histograms shown are normalized to the modal value.

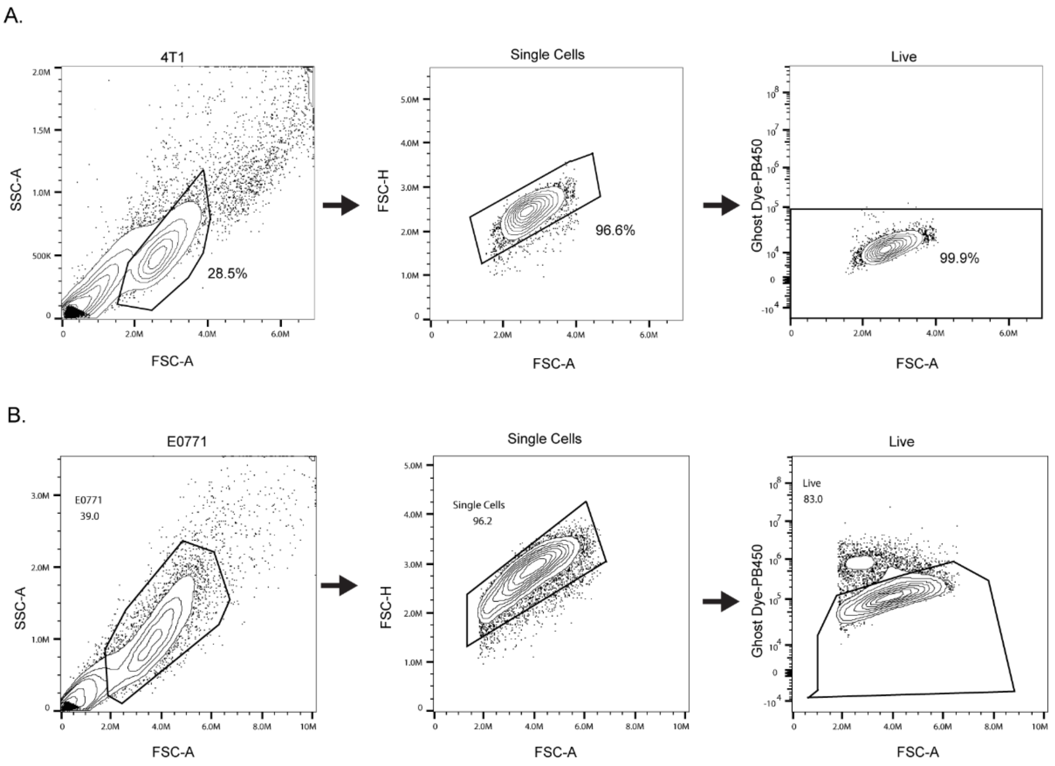


**Supplementary Figure 5. Gating strategy of 4T1 and E0771 cells for staining ligands of NK cell receptors.** 4T1 cells (A) and E0771 cells (B) that were gated negatively for Ghost Dye (live cells) were analyzed for expression of ligands for NKG2D and DNAM-1 as seen in Figure 3.

Ben-Shmuel et al. Supplementary Figure 6

A.

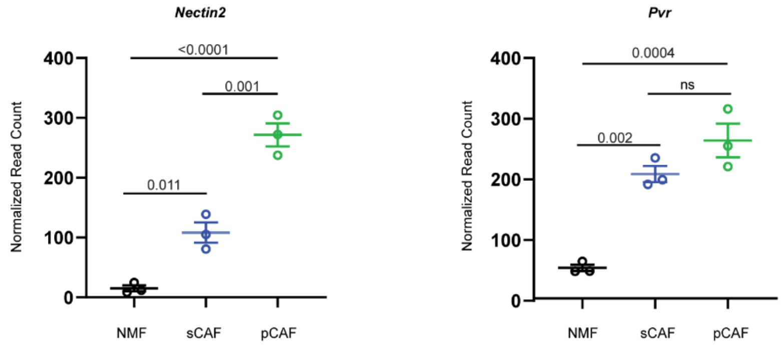


B.

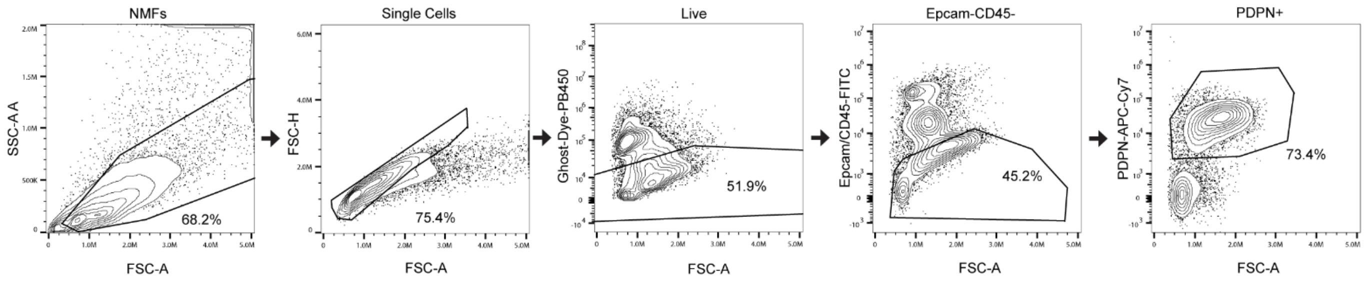


C.

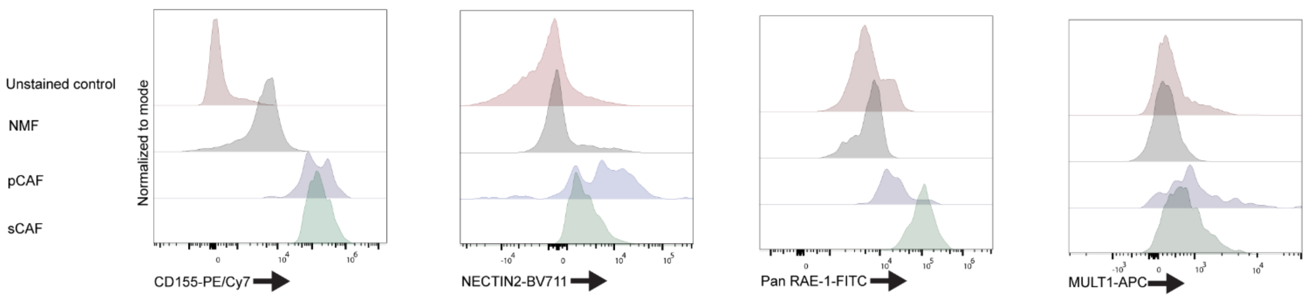


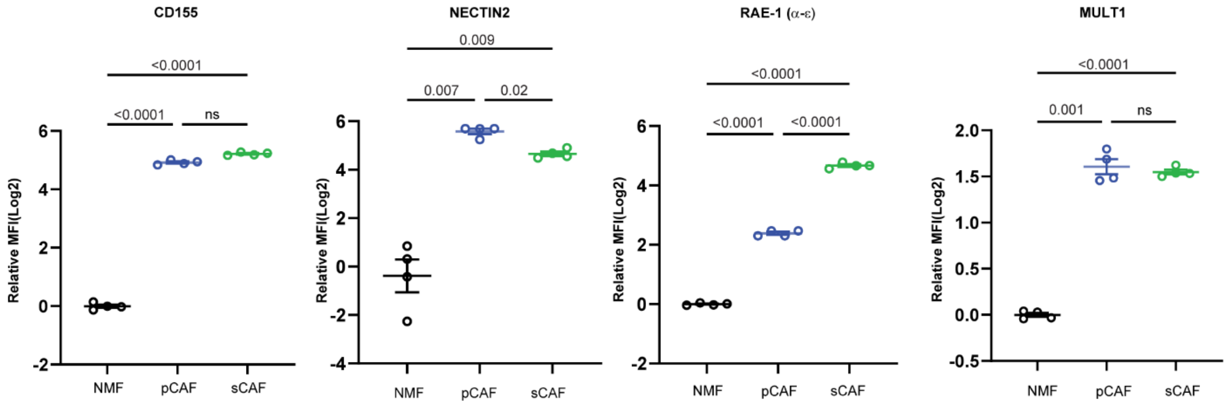

**Supplementary Figure 6. Upregulation of ligands for NK cell receptors on CAFs.** (A) Differentially expressed gene quantification of ligands for the DNAM-1 receptor, *Nectin2* and *Pvr* on NMFs, pCAFs, and sCAFs freshly isolated from fat pads or 4T1 tumors of BALB/C mice. Data are displayed as mean ± SEM of N=3 biologically independent experiments. (B) Gating strategy for FACS analysis of NMFs freshly isolated from murine fat pads. Live NMFs were negatively selected for epithelial (EPCAM) and immune (CD45) markers, and positively selected for podoplanin staining (PDPN). ( (C) FACS quantification of ligands for NKG2D and DNAM-1 expressed on NMFs, pCAFS, and sCAFs freshly isolated from fat pads or E0771 tumors of C57BL/6 mice. MFI of all live cells are normalized to NMF samples and log2 transformed. Data are presented as mean ± SEM of N=4 biologically independent experiments. FACS histograms are normalized to the modal value.

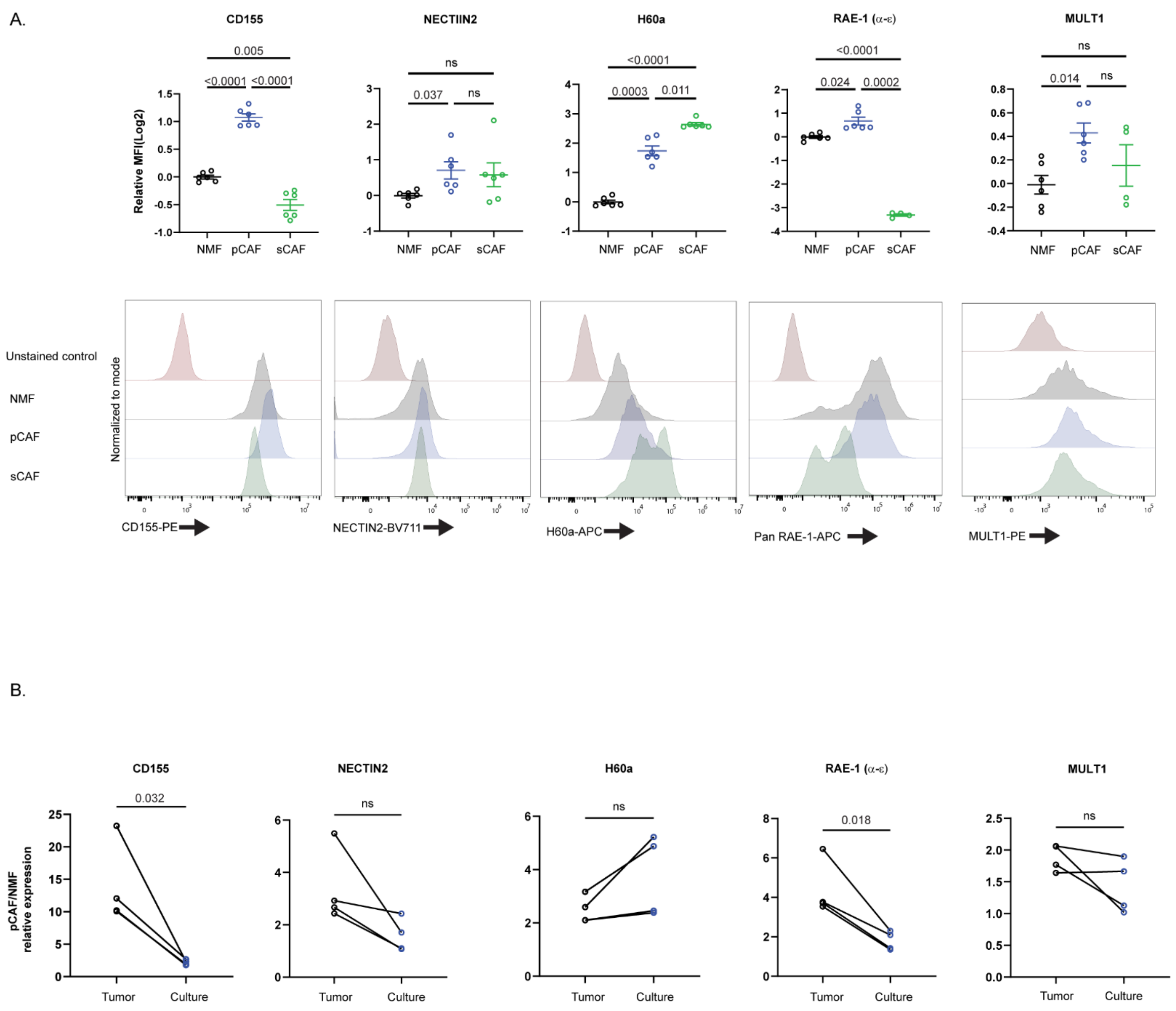


**Supplementary Figure 7. NMFs upregulate NK ligands following ex-vivo culture.** (A) FACS quantification of ligands for NKG2D and DNAM-1 expressed on NMFs, pCAFs, and sCAFs isolated from fat-pads or 4T1 tumors of BALB/C mice following ex-vivo culture for 4 days. MFI of all live cells are normalized to NMF samples and log2 transformed. Data are presented as mean ± SEM of N=6 biologically independent experiments. FACS histograms are normalized to the modal value. (B) Comparison of fold change of ligand expression between CAFs and NMFs freshly isolated from tumors or following ex-vivo culture for 4 days. The ratio between expression of NK ligands on pCAFs and NMFs is shown for experiments conducted on cells fresh from tumors or following ex vivo culture for 4 days. Comparisons were conducted for N=4 biologically independent experiments.

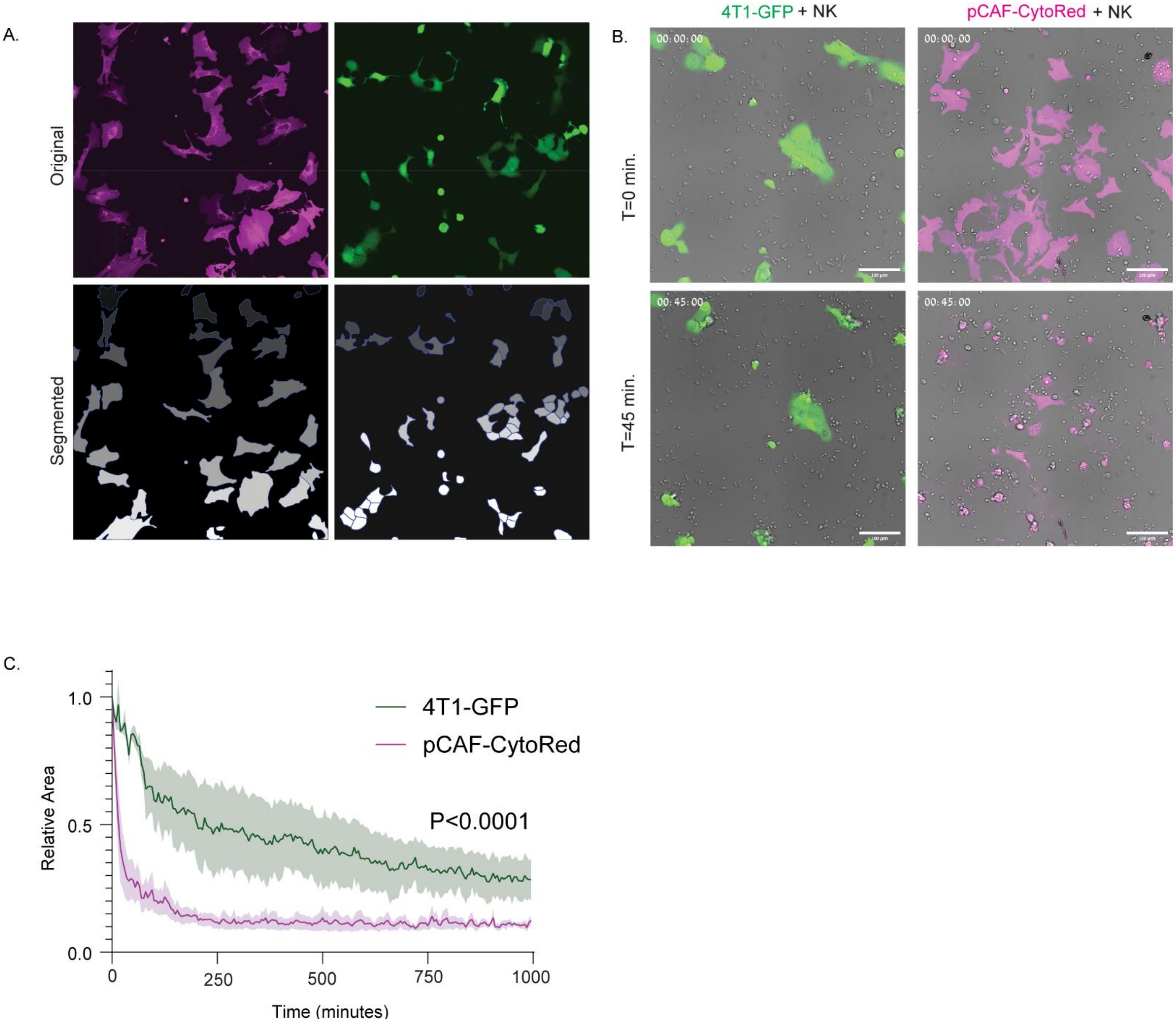


**Supplementary Figure 8. CAFs are susceptible to NK cell-mediated lysis** (A) Segmentation of 4T1-GFP and pCAFs as described in the materials and methods. (B+C) Representative image and quantification of 4T1-GFP cells or pCAF lysis by NK cells in co cultures as a function of time. Average cell area of either 4T1 or pCAF labeled and segmented cells was measured for each time point (see materials and methods). The average cell area for each time point was normalized relative to the average cell area during time point 0. The graph represents 3 independent biological experiments, and displays the relative average cell area of 4T1 or pCAF cells for each time point, ± SEM. Statistical analysis was conducted using two-way ANOVA with Time and cell type as discrete factors. The P value is provided for Time * Cell type interaction.

Ben-Shmuel et al. Supplementary Figure 9

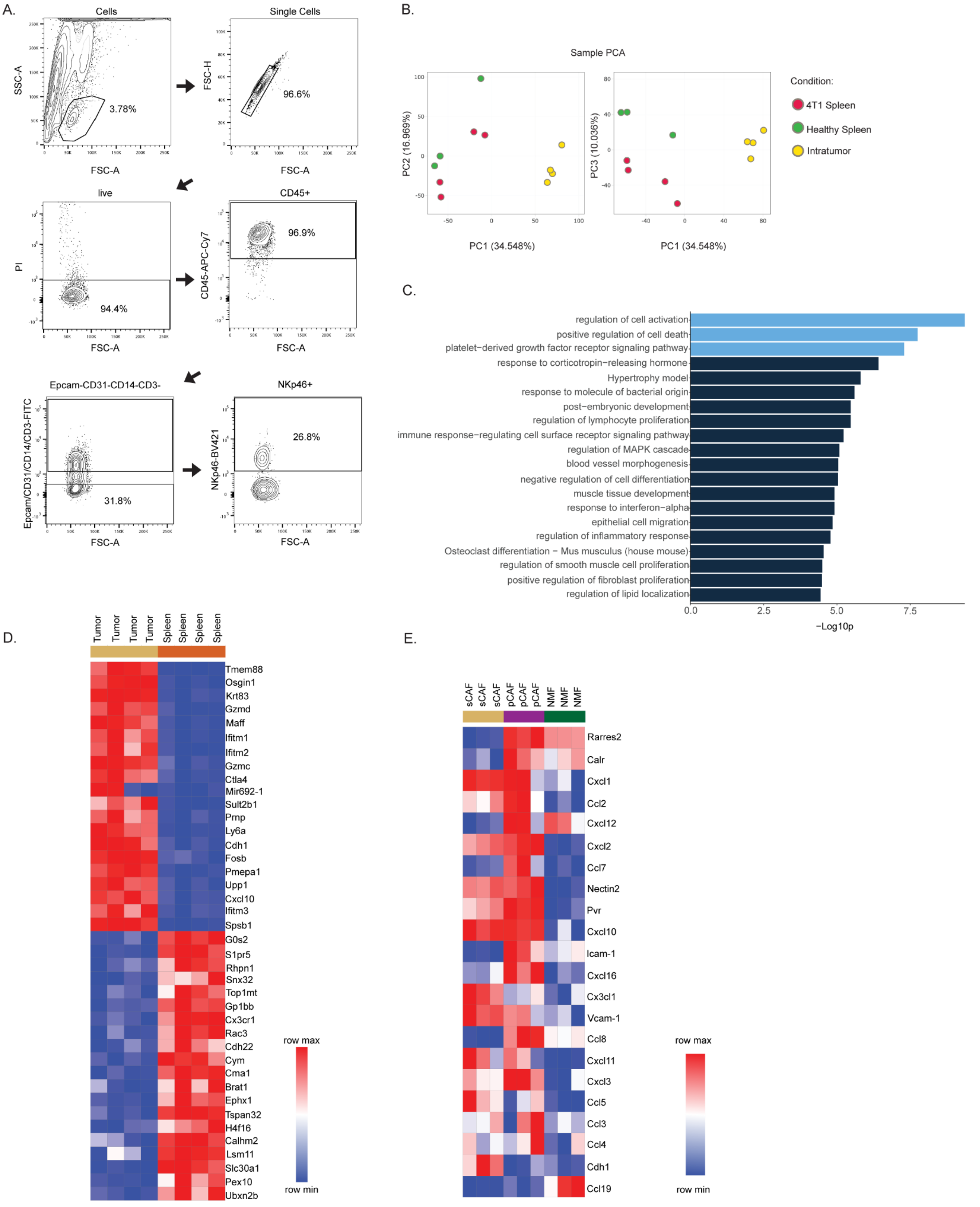

**Supplementary Figure 9. Tumor infiltrating NK cells are characterized by a dysfunctional phenotype and may be recruited to CAFs via chemotactic signals** (A) Gating strategy for isolation of NK cells from 4T1 tumors. (B) PCA analysis of bulk RNA-seq of splenic NK cells from healthy and 4T1-bearing mice and 4T1 infiltrating NK cells. (C) Pathway analysis of the top 100 genes upregulated in tumor infiltrating NK cells compared to splenic NK cells from 4T1-bearing mice. (D) Heatmap of top differentially expressed genes between tumor infiltrating NK cells compared to splenic NK cells from 4T1-bearing mice. (E) Heatmap of gene expression of ligands for NK cell activating, adhesion, and chemotactic receptors by sCAFs and pCAFs from 4T1 tumors or NMFs from healthy mice.

Ben-Shmuel et al. Supplementary Figure 10

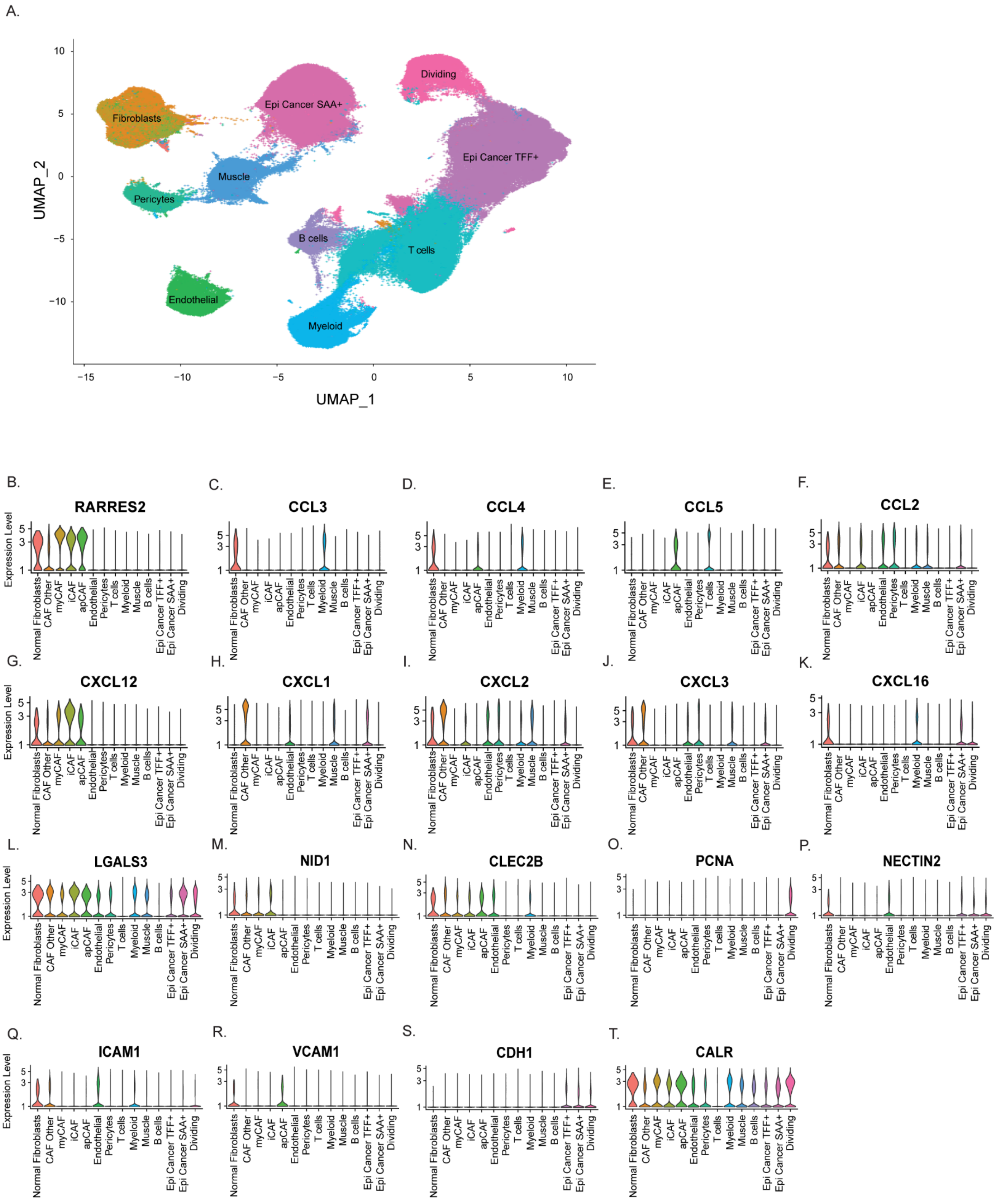

**Supplementary Figure 10. Comparison of transcript expression for NK cell chemotactic, adhesion, and activating receptors in human TNBC tumors** (A-B) Single-cell RNA-seq data of breast cancer tumors was reanalyzed using the Seurat R toolkit. (A) Integrated Uniform Manifold Approximation and Projection (UMAP) of 62 patients over 343811 cells. Overall, 10 distinct clusters were observed: dividing cancer cells, SAA+ and TFF+ cancer cells, T-cells, B-cells Myeloid cells, Endothelial cells, pericytes, fibroblasts and muscle cells. 5 fibroblasts subclusters were annotated: normal, iCAF, myCAF, apCAF, and undefined (other). (B) Violin plots of normalized expression levels of ligands for NK cell receptors expressed in each cluster.  Violin plots shown here were selected for genes that were highly expressed in at least 1 cluster.

Ben-Shmuel et al. Supplementary Figure 11

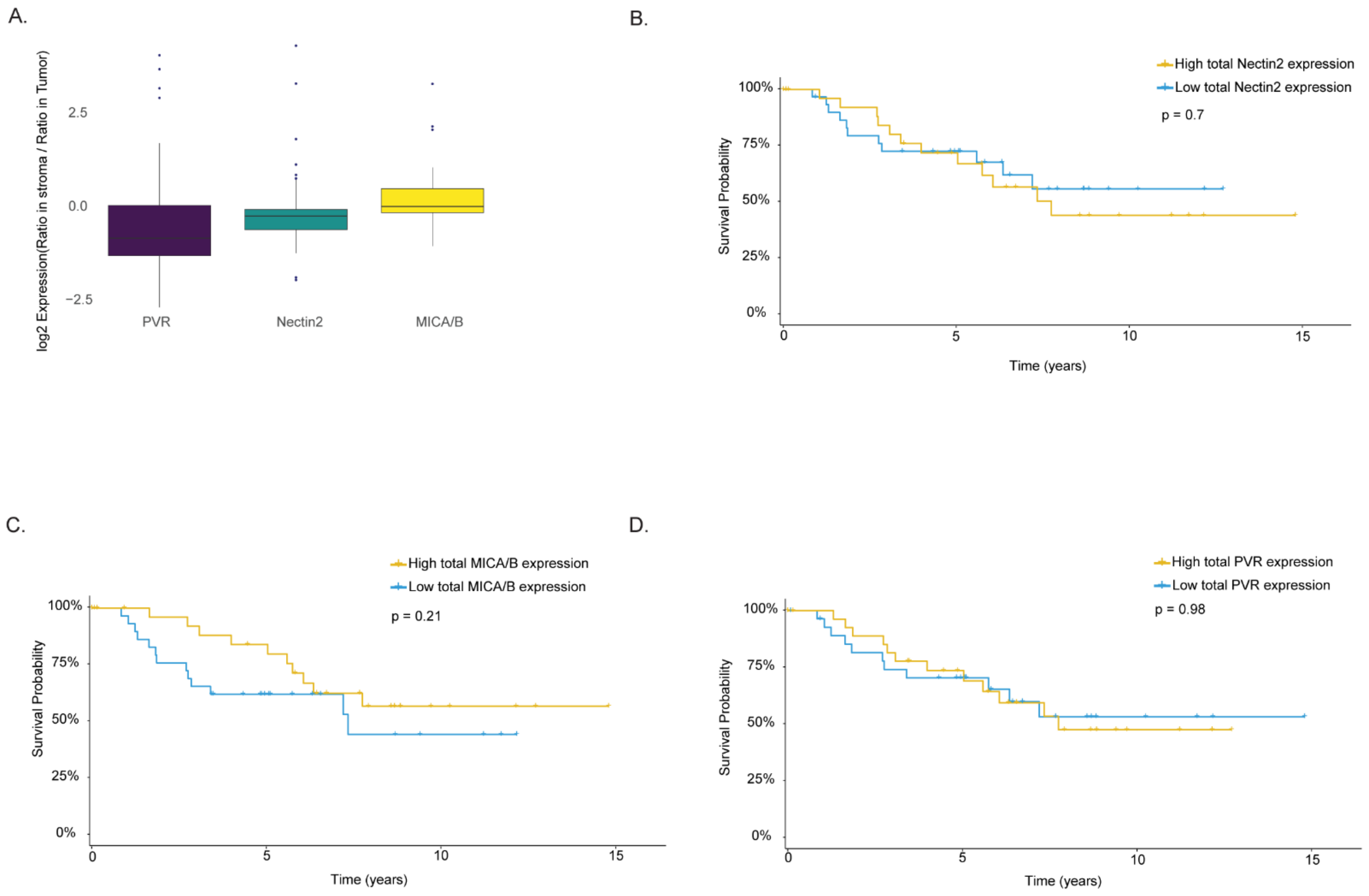


**Supplementary Figure 11. Association of NK ligand expression on CAFs and patient outcomes in TNBC.** (A) Relative expression of staining of PVR, NECTIN2, and MICA/B in patient TMAs between non immune stroma cells (CK-CD45-) and cancer cells (CK+). Data is presented as Log2 expression of stroma/cancer in tumors. (B-D) Kaplan Meier survival analysis of patients stratified according to low (below median) or high (above median) NECTIN2, MICA/B, or PVR staining of total tumor area.
