## supplementary videos legend for "Cancer-associated fibroblasts serve as decoys to suppress NK cell anti-cancer cytotoxicity"

**Supplementary Video 1**

4T1-GFP cells (green) and cytopainter red-labeled pCAFs (magenta) were seeded as described in Methods. NK cells from spleens of BALB/C mice were activated overnight with IL-2+IL-15 and were introduced to wells with 4T1+pCAFs and immediately imaged for 16 hours. This is video is from 1 of 3 biologically independent experiments.

**Supplementary Video 2**

4T1-GFP cells (green) and cytopainter red-labeled pCAFs (magenta) were seeded as described in Methods. NK cells from spleens of BALB/C mice were activated overnight with IL-2+IL-15 and were introduced to wells with 4T1+pCAFs and immediately imaged for 16 hours. This is video is from a 2nd of 3 biologically independent experiments.

**Supplementary Video 3**

4T1-GFP cells (green) and cytopainter red-labeled pCAFs (magenta) were seeded as described in Methods. NK cells from spleens of BALB/C mice were activated overnight with IL-2+IL-15 and were introduced to wells with 4T1+pCAFs and immediately imaged for 16 hours. This is video is a close-up ROI of Supplementary Video 2.

**Supplementary Video 4**

4T1-GFP cells were seeded as described in Methods. NK cells from spleens of BALB/C mice were activated overnight with IL-2+IL-15 and were introduced to wells with 4T1 cells and immediately imaged for 16 hours.

**Supplementary Video 5**

Cytopainter red-labeled pCAFs were seeded as described in Methods. NK cells from spleens of BALB/C mice were activated overnight with IL-2+IL-15 and were introduced to wells with 4T1 cells and immediately imaged for 16 hours.
